# Defense mechanism of a bacterial retron supramolecular assembly

**DOI:** 10.1101/2023.08.16.553469

**Authors:** Yanjing Wang, Chen Wang, Zeyuan Guan, Jie Cao, Jia Xu, Shuangshuang Wang, Yongqing Cui, Qiang Wang, Yibei Chen, Delin Zhang, Ming Sun, Pan Tao, Tingting Zou

**Author notes:** **Correspondence:** Tingting Zou, Pan Tao.

## Abstract

Retrons are a class of multigene antiphage defense system typically consisting of a retron reverse transcriptase, a non-coding RNA, and a cognate effector. Although the triggers for several retron systems have been discovered recently, the full picture of how retron systems sense invading phages and mediate defense remains to be elucidated. Here, we focus on the retron Ec86 defense system and report its modes of activation and action. We identified a phage-encoded DNA cytosine methyltransferase (Dcm) as the trigger of the Ec86 system and show that Ec86 senses msDNA methylation and becomes activated. We further determined the structure of a tripartite retron Ec86 supramolecular assembly, which is primed for activation by Dcm, and demonstrated that the activated system confers defense through depletion of nucleoside derivatives. These findings emphasize the role of retrons being a second line of defense and highlight an emerging theme of anti-phage defense through supramolecular complex assemblies.

## Introduction

During the arms race between bacteria and their phage foes, bacteria have evolved a broad range of molecular defense systems that involve various enzymatic activities against phage infections^1–6^. The most abundant and widespread antiphage defense systems restriction-modification (RM) and CRISPR-Cas, which have long been recognized as major lines of bacterial defense, sense and degrade invading nucleic acids^1,7–10^. Continued investigation of the bacteria-phage warfare has expanded the bacterial antiphage defense system repertoire, with dozens of novel systems being discovered in the past few years^1,3,5,11–14^. Recent identification of phage components that trigger different defense systems indicates that diverse mechanisms of activation exist^11,15–20^. In the meantime, advances in the determinations of the defense mechanisms underpinning several systems revealed that activated immune effectors have various modes of action, including disrupting host cell membrane integrity, depleting molecules essential for phage replication, and producing small molecules that block phage replication^11,21–32^. Despite this, the defense mechanisms of most novel defense systems remain largely uncharacterized.

Retrons are an intriguing class of multigene defense systems that typically comprises three components: a retron reverse transcriptase (RT), a non-coding RNA (ncRNA), and a cognate effector^3,11,33,34^. The ncRNA comprises *msd* and *msr*, of which the *msd* part is reverse-transcribed into msdDNA by retron RT, eventually resulting in the production of a chimeric RNA-DNA molecule called the multicopy single-stranded DNA (msDNA)^11,35–39^. The effector encodes an accessory protein or RT-fused domain, which usually has enzymatic functions crucial for antiphage defense^3,11,40^. A recent study has suggested that the three retron components together form a tripartite toxin-antitoxin (TA) system, in which RT and msDNA form the antitoxin unit that neutralizes the effector’s toxicity^33^. Corroborating this proposal, the effector of retron Ec86 (also known as Eco1, the first retron discovered in *Escherichia coli*) has been shown to stably associate with the mature Ec86 (RT-msDNA) and form an RT-msDNA-effector complex^34^. These advances have deepened our understanding of retrons, but the full picture of how retron systems recognize invading phages and exert antiphage functions remains to be elucidated.

Here, we focused on the retron Ec86 defense system from *E. coli* BL21(DE3)^11^, and investigated its modes of activation and action (Figure 1A). We identified a phage-encoded DNA cytosine methyltransferase (Dcm) as the trigger of the Ec86 system and demonstrated that Ec86 msDNA senses Dcm-mediated methylation in its DNA stem loop (DSL) region. We further determined the cryo-electron microscopy (cryo-EM) structure of a tripartite retron Ec86 supramolecular assembly, namely the Ec86-effector filament. Combined with in vivo and in vitro biochemical analyses, we demonstrated that the Ec86-effector filament represents a fully assembled Ec86 defense system primed for activation and the activated system confers phage defense through depleting nucleoside derivatives. These highlight a mechanism of antiphage defense through sensing of phage-encoded proteins and functioning via depletion of molecules essential for phage replication.

**Figure 1.**
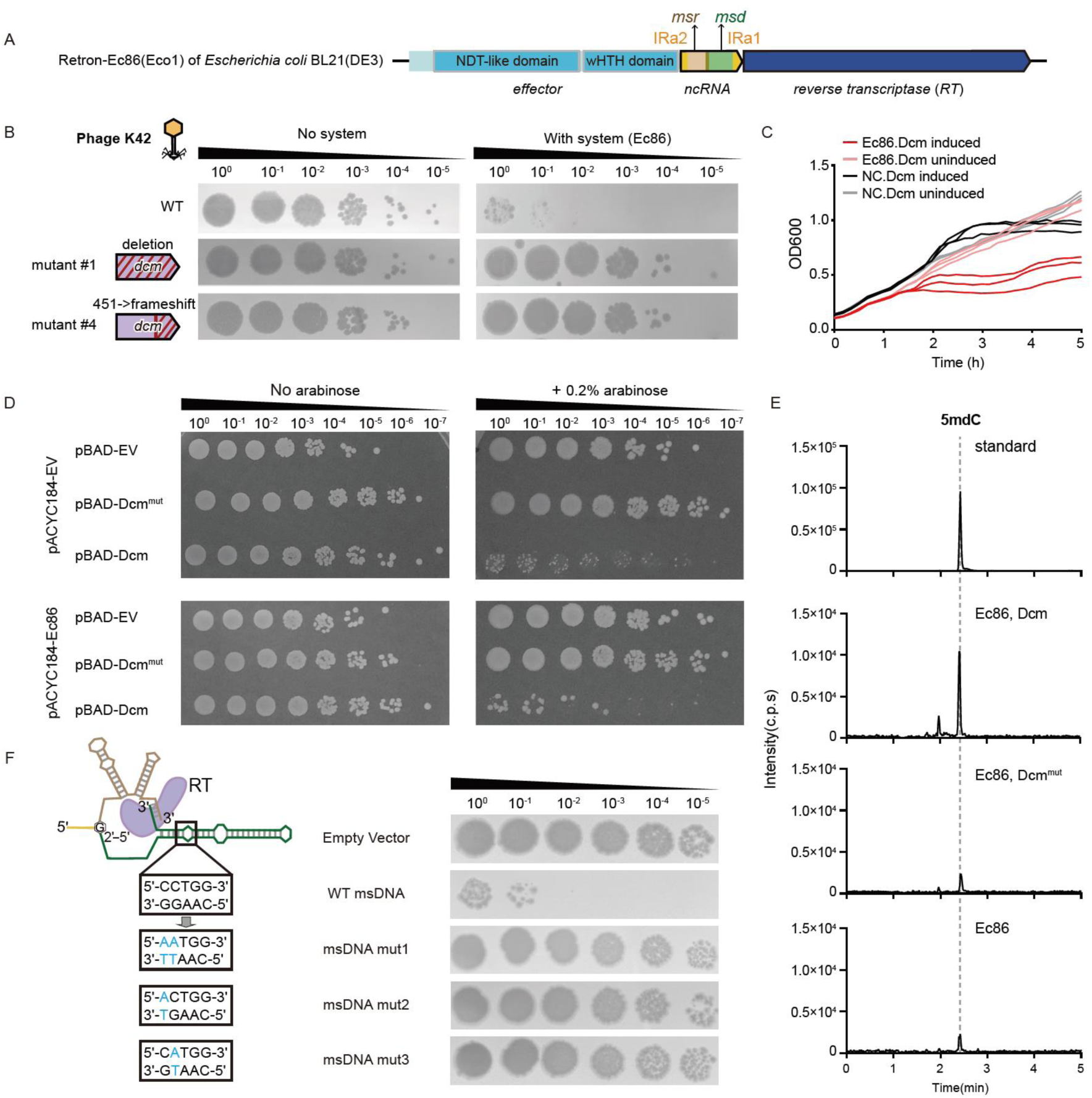
Retron Ec86 senses msDNA methylation by a phage-encoded DNA cytosine methyltransferase (Dcm). (**A**) Schematic of the retron Ec86 system (retron Ec86 gene cassette and its cognate effector) in the E. coli BL21(DE3) genome. (**B**) Representative phage mutants capable of escaping the retron Ec86 system defense. Shown are 10-fold serial dilution plaque assays for the WT and mutant phages on the *E. coli* MG1655 strain transformed with an empty vector or a plasmid containing the WT Ec86 system. Images are representative of three replicates. The mutations in mutant phages are presented on the right. (**C**) Growth curves of *E. coli* MG1655 strain expressing the Ec86 system or a no-system control (NC) plasmid that lacks the system in the presence and absence of Dcm expression. The expression of the phage-encoded Dcm protein was induced by the addition of 0.2% arabinose. Three biological replicates are presented as individual curves, each a mean of two technical replicates. The experiment was repeated twice. (**D**) Serial dilutions of *E. coli* MG1655 cells expressing the retron Ec86 system from its native promoter or empty vector and indicated Dcm, Dcm^mut^ (that contains C82A, E128A, K175A, and K176A mutations), or the corresponding empty vector from an arabinose-inducible promoter on LB media containing no or 0.2% arabinose. Images are representative of three replicates. (**E**) Liquid chromatography–mass spectrometry (LC-MS) analysis of C5 cytosine methylation of Ec86 msDNA. The msDNA was purified from *E. coli* MG1655 strains co-expressing Ec86-effector defense system and Dcm, Dcm^mut^ (that contains C82A, E128A, K175A, and K176A mutations) or empty vector. Top panel, synthetic 5mC standard control. (**F**) Mutational analysis of msDNA in the 5′-C**<u>C</u>**WGG-3′ Dcm methylation site. Shown are 10-fold serial dilution plaque assays for the WT phage on the *E. coli* MG1655 strain transformed with WT or mutated Ec86 systems. Images are representative of two replicates.

## Results

### Phage-encoded Dcm-like C5 cytosine methyltransferase triggers retron Ec86

We sought to understand how the retron Ec86 system senses invading phages through isolating phage mutants that escape Ec86 defense (escaper phages). We hypothesize that the escapers harbor mutations in phage components that activate the defense system and thus could be potential candidates directly recognized by retron Ec86 defense system. We selected the phage KarlJaspers strain Hzau42 from the Hzau phage collection (hereafter referred to as phage K42), whose genome (∼51.7 kb) displays a pairwise identity of 95.4% with that of phage KarlJaspers strain Bas06 (∼51.5 kb), for the generation of escaper phages. Four phage K42 escaper mutants were isolated and sequenced. We identified the mutations in each of the escaper phages using the wildtype (WT) K42 phage genome as the reference. Notably, all four escapers (mut#1-#4) harbor mutations that disrupt (or potentially disrupt) the 711-bp gene encoding a DNA cytosine methyltransferase (hereafter called *dcm* and Dcm for the gene and encoded protein, respectively). Specifically, mut#1 lost a 2,262-bp region that contains four genes including *dcm* in its genome, and mut#4 showed a 1-bp indel (deletion of dG451) in the *dcm* gene which resulted in a frameshift with a premature stop codon (Figure 1B). Both K42 mut#2 and mut#3 exhibit a single-base mutation (dC-to-dA) 9-bp upstream of the start codon of *dcm*. These suggest that mutations disrupting the expression of Dcm may have enabled these K42 mutants to overcome the Ec86 defense system.

To investigate whether Dcm is the phage component that triggers the defense system, we expressed Dcm derived from phage K42 in *E. coli* MG1655 and checked their growth in liquid culture and on plates. Following the expression of Dcm, a marked decrease in growth was observed in bacteria that contain the Ec86 retron defense system, but not in bacteria lacking the system (Figure 1C). We also identified the putative catalytic residue (cysteine 82, C82) and substrates binding residues (E128, K175, and K176) of phage K42 Dcm through AlphaFold2 predictions and subsequent structural alignment with the MTase domain of DNA methyltransferases 3A (DNMT3A, PDB: 6F57)^41^ (Figures S1A and S1B). Thus, we generated a Dcm^mut^ construct that contains alanine substitutions of these residues (i.e., C82A, E128A, K175A, and K176A) and included it in plate assays. Corroborating the observations in liquid culture, a 3-fold decrease was observed in the efficiency of plating (EOP) when overexpressing Dcm in cells containing the Ec86 defense system (Figure 1D). In addition, we also examined the colony forming units (CFU) of *E. coli* BL21(DE3), a common laboratory strain of *E. coli* that contains a native retron Ec86 operon, and the corresponding *E. coli* BL21(DE3) Δ*Ec86* strain after transformation of plasmids that contain inducible Dcm or Dcm^mut^. While both plasmids were transformed with similar efficiency into both *E. coli* strains in the absence of arabinose, the transformation of the plasmid containing Dcm into *E. coli* BL21(DE3) consistently and repeatedly failed in the presence of arabinose, yielding zero colonies or a single colony in three independent transformation attempts (Figure S1C). Furthermore, we sequenced all WT-Ec86-sensitive phages from the Hzau collection and found that of the 35 phages from *Drexlerviridae*, *Siphoviridae*, and *Demerecviridae* families, 34 encode Dcm (Figure S2). These results together indicate that Dcm is the trigger of the Ec86 defense systems.

### Dcm methylates the stem loop region of msDNA to activate retron Ec86

As Dcm is predicted to have the capacity to directly interact with and methylate double-stranded DNA (dsDNA), we examined its methylation activity toward the dsDNA hairpin part, known as the DNA stem loop (DSL) region, of Ec86 msDNA. We transformed Dcm, Dcm^mut^, or empty vector control, respectively, into *E. coli* MG1655 cells containing the Ec86 defense system, purified the msDNA from corresponding samples and checked their C5 cytosine methylation levels via liquid-chromatography followed by mass spectrometry (LC-MS). The Dcm sample exhibited a high C5 cytosine methylation level, while low C5 cytosine methylation levels were detected for the msDNA from the Dcm^mut^ and empty vector control samples (Figures 1E). This suggests that Dcm can methylate Ec86 msDNA.

Considering that Dcm methylates 5′-C**<u>C</u>**WGG-3′ (W stands for A or T; the bold underlined **<u>C</u>** is methylated) in dsDNA^42^ and the DSL region of Ec86 msDNA contains a Dcm motif (5′-C**<u>C</u>**TGG-3′), we wondered if the methylation of the Dcm motif is responsible for activation. We generated three mutated Ec86 systems, in which the Dcm motif 5′-C**<u>C</u>**TGG-3′ in DSL was replaced with 5′-AATGG-3′, 5′-ACTGG-3′, or 5′-CATGG-3′ without disrupting the hairpin structure, and subjected them to antiphage assays against phage K42. These mutations abolished phage defense activity (Figure 1F), corroborating the importance of this methylation site in msDNA. These results together suggest that the identified phage K42 Dcm methylates the 5′-C**<u>C</u>**TGG-3′ motif in the DSL region of msDNA and activates the Ec86 defense system.

### Retron Ec86 and its effector form a supramolecular filament assembly

To further understand the mechanistic basis of anti-phage defense by retron Ec86 and its cognate effector after trigger identification, we attempted to obtain the active state Ec86-effector and/or effector proteins by co-expressing the defense system with its trigger. However, very few *E. coli* BL21 cells can be harvested after the co-transformation of *dcm*- and Ec86-system-containing plasmids and IPTG induction, likely due to the toxicity of Dcm itself and the activated Ec86 system. On the other hand, we noticed that only one peak (at ∼14 ml) with a homogeneous Ec86 (RT-msDNA) complex can be obtained during size-exclusion chromatography (SEC), while several higher-order multimeric peaks (at ∼11.5 ml, ∼12.5 ml, and ∼14 ml) were observed in SEC during the purification of the effector-bound Ec86 (Figure S3). This indicates the formation of higher-order Ec86-effector ternary complexes. The fractions (at ∼11.5 ml) corresponding to the higher molecular weight species were used for negative stain electron microscopy, cryo-EM grid screening and subsequent structural analyses. Visualization of the higher molecular weight species revealed that retron Ec86 and its effector formed supramolecular filament (Figure 2A; Figure S3). We further solved the structure of the Ec86-effector filament assembly at 2.7 Å (Figure 2B; Figure S3). In this supramolecular filament assembly, a tetrameric Ec86-effector unit (with the RT, msDNA, and effector stoichiometric ratio of 4:4:4) consisting of four RT-msDNA-effector protomers defines the minimal structural repeating unit (Figure 2C). The tetrameric unit measures ∼320 Å in length and contains all interaction interfaces needed for stable filamentous packing. The units stack head-to-tail and twist (with a helical twist of approximately ∼75°) alongside the longitudinal axis lengthwise, forming supramolecular filaments.

**Figure 2.**
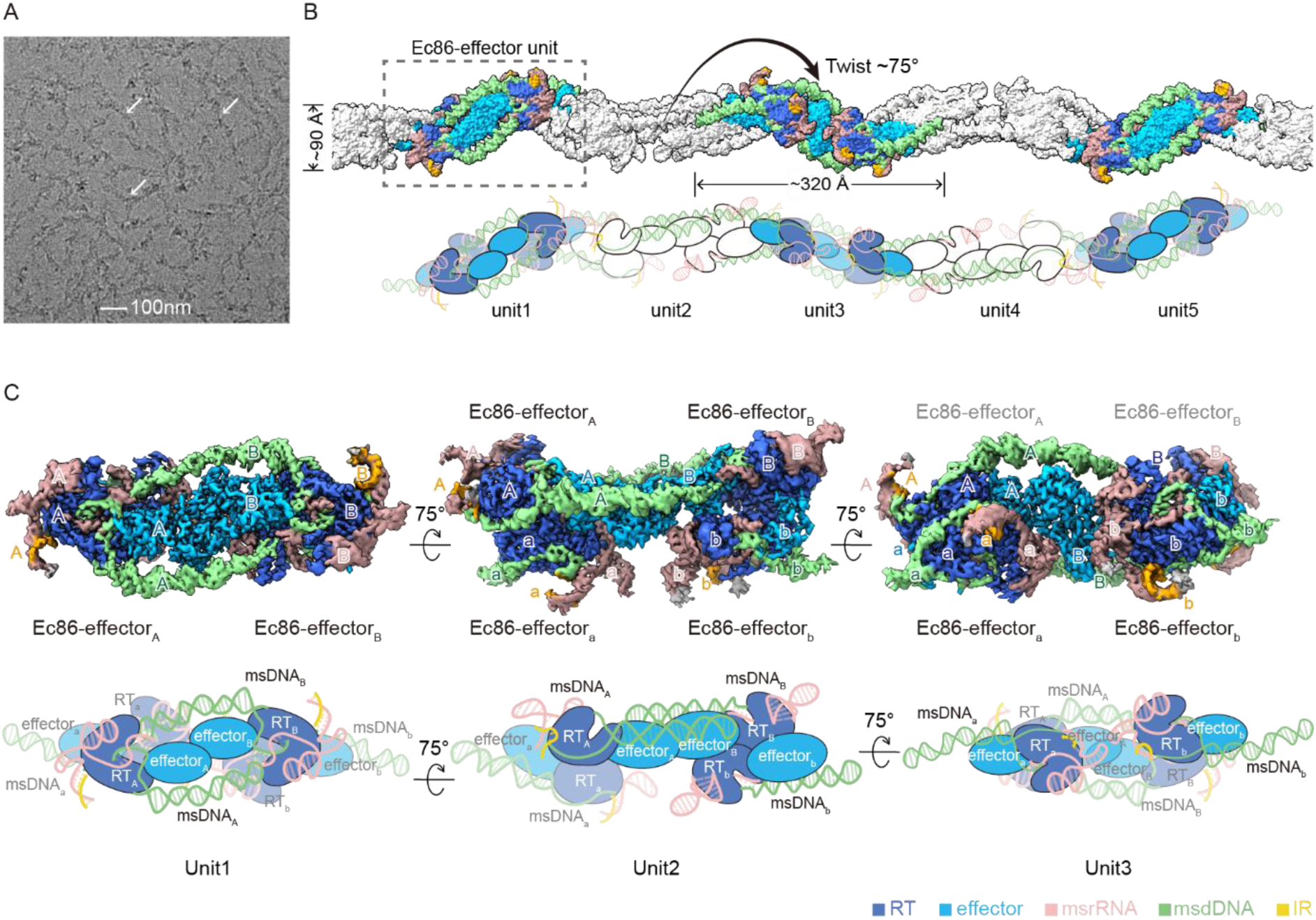
Overall structure of the Ec86-effector filament and its minimal structural repeating unit. (**A**) A cryo-EM micrograph of retron Ec86-effector filament assemblies. Scale bar, 100 nm. (**B**) Schematic representation of the Ec86-effector filament complex. The gray dashed box highlights an isolated tetrameric Ec86-effector repeating unit. msrRNA, rosy brown; msdDNA, light green; reverse transcriptase, royal blue; IRs (inverted repeats), orange; effector, deep sky blue. (**C**) A close-up view of the tetrameric Ec86-effector repeating unit. Top, cryo-EM density map of the repeating unit. Bottom, schematic representation of the repeating unit.

A closer inspection of the structure revealed that the four protomers within one tetrameric repeating unit can be further grouped into two subunits, each exhibiting high similarity with the previously reported dimeric effector-bound Ec86 complex (Figure S4)^34^. For clarity, we name the four protomers in the order of “aABb” and the two dimeric complexes subunit_A_ (comprising protomers_a/A_) and subunit_B_ (comprising protomers_b/B_) (Figure 2C). In each subunit, two reverse transcriptases wrapped by msDNA form a scaffold, and two effectors fill the space in between, forming the same msDNA-RT, msDNA-effector, and RT-effector interaction interfaces found in the effector-bound Ec86 complex (Figure S4). Differences were also identified between the subunits and the effector-bound Ec86 in that the observed msDNAs are intact and the bilobed effectors exhibit a more compact conformation in subunit_A_ and subunit_B_ (Figure 2C; Figures S4 and S5). The two subunits are further assembled into the tetrameric repeating unit via direct msDNA-effector and effector-effector contacts between protomer_A_ and protomer_B_ from subunit_A_ and subunit_B_, respectively, resulting in the “sandwiching” of effector_A_ and effector_B_ between the two DNA stem loop regions (DSLs, DSL_A_ and DSL_B_) (Figure 2C). The same subunit_A_-subunit_B_ interaction modes were observed within and between neighboring repeating units, contributing to stable Ec86-effector filamentous packing.

### Ec86 msDNA DSL directly contacts cognate effector

As mentioned above, the interactions within each of the two subunits in the tetrameric repeating unit have been described before^34^, we thus focused on the newly observed intra- and inter-subunit interaction interfaces within and between subunit_A_ and subunit_B_. For the intra-subunit msDNA-effector interaction, the msDNA DSL region binds the positively charged surface areas of the effector mainly through electrostatic interactions (Figure 3A). The bilobated Ec86 effector consists of 14 α-helices and 7 β-strands. Taking protomer_B_ as an example, the effector N-lobe nucleoside deoxyribosyltransferase-like (NDT-like) domain (α2–9 and β1–5) contains a large region rich in positively charged residues that face the DSL part of msDNA (Figure 3A; Figure S6A). This effector_B_ positively charged region attracts and interacts with the msDNA_B_ DSL, forming an extensive electrostatic interaction network. Moreover, the lysine residues located in and close to α9 were also observed to interact with the major groove of msDNA_B_ DSL through hydrogen bond interactions. Specifically, residues K149 (in the loop between α7 and β5), K177, K178, and K185 (in α9), and K29 (located in the loop connecting α1 and β1) interact with the backbones of dG55, dT56, dT57, and dT25 in the msDNA_B_ DSL (Figure 3B). Individual K-to-A mutations (K29A, K149A, K177A, and K185A, except for K178A) abrogated the antiphage activity of the Ec86 defense system, while *E. coli* MG1655 transformed with the WT Ec86 defense unit exhibited defense activity against phage (Figure 3C). For the inter-subunit msDNA-effector interaction between protomer_A_ and protomer_B_. Taking msDNA_B_ and effector_A_ as an example, msDNA_B_ directly contacts the C-lobe of effector_A_ (Figure 3D). Hydrogen bonds were observed between effector_A_ R219 and msDNA_B_ dC44/dG45. Two-base-pair and four-base-pair deletions of the bases between msDNA DSL loop1 and loop2 showed little impact on phage defense (Figure 3E). Taken together, these results indicate that the intra-subunit (rather than inter-subunit) effector-msDNA interactions play important roles in antiphage activity.

**Figure 3.**
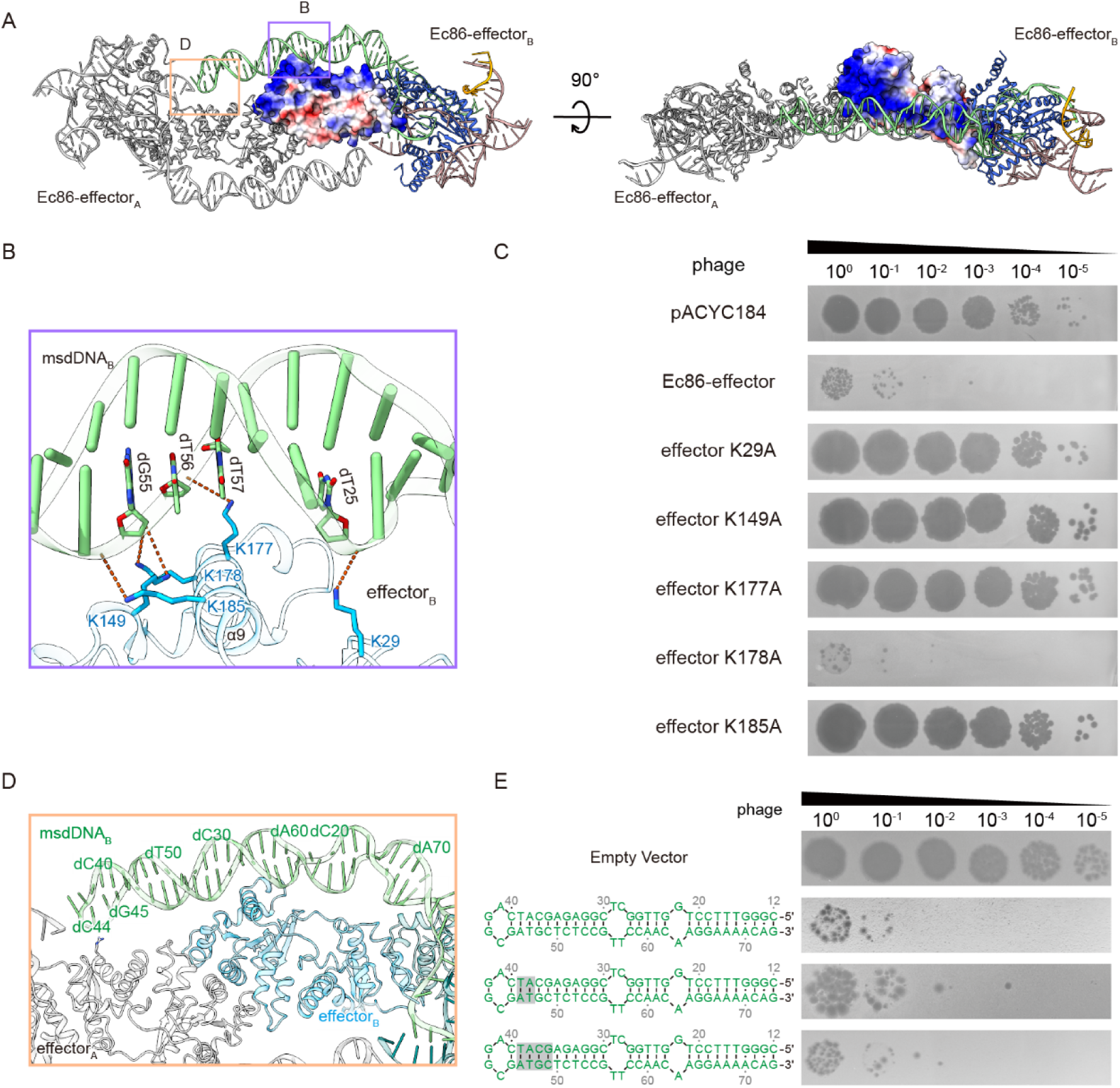
The msDNA DNA stem loop (DSL) region directly interacts with the effector NDT-like domain. (**A**) Analysis of msDNA-effector interaction interfaces based on the cryo-EM model. The msDNA DNA stem-loop (DSL) region is bound to the positively charged surface areas of the two effectors through electrostatic interactions. The two effectors are shown as cartoon model and electrostatic surface potential model, respectively. The msDNAs are shown in cartoon. (**B**) A close-up view of the indicated interaction interface between msDNA_B_ DSL and effector_B_. (**C**) Serial dilution plaque assays shown for the *E. coli* MG1655 strain transformed with plasmids encoding WT or mutated Ec86 systems. Images are representative of two replicates. (**D**) A close-up view of the indicated interaction interface between msDNA_B_ and effector_A_. (**E**) Mutational analysis of msDNA DSL within Ec86-effector system. Shown are 10-fold serial dilution plaque assays for the *E. coli* MG1655 strain transformed with plasmids encoding WT or mutated Ec86 systems with truncated versions of msDNA. The truncated regions in msDNA are indicated by gray shadows. Images are representative of two replicates.

### Ec86 effector dimerization is indispensable for antiphage defense

Apart from the msDNA-effector interaction, inter-subunit effector-effector interaction is also observed between protomer_A_ and protomer_B_ in the repeating unit. Specifically, effector_A_ is bound to effector_B_ via a composite interface involving two NDT-like domains, forming a tight homodimer with an asymmetric interaction interface (Figure 4A). Extensive hydrophobic packing interactions and a network of hydrogen bond interactions were found between the two NDT-like domains, similar to those observed in conventional deoxyribosyltransferases. Hydrogen bond interactions can be observed between effector_A_ α5 and effector_B_ α3-5 and α4, resulting in the close packing of the α5s from both effectors. The side chains of effector_A_ N112 and R141 form hydrogen bonds with the main chains of effector_B_ A109 and L72/Q75, respectively. Side-chain hydrogen bond interactions were also observed between effector_A_ N112 and N113 with effector_B_ E84 and N112 (Figure 4B; Figure S6B). Individual alanine substitutions (E84A, N112A, N113A, and R141A) in the observed interface strongly reduced or abolished antiphage activity (Figure 4C). In addition, the Ec86 system containing the effector^N112A^ mutant eluted mainly at the position of the low molecular fractions in gel filtration. The systems containing the individual effector^E84A^, effector^N113A^, and effector^R141A^ mutants showed elution profiles similar to the WT system (Figure 4D).

**Figure 4.**
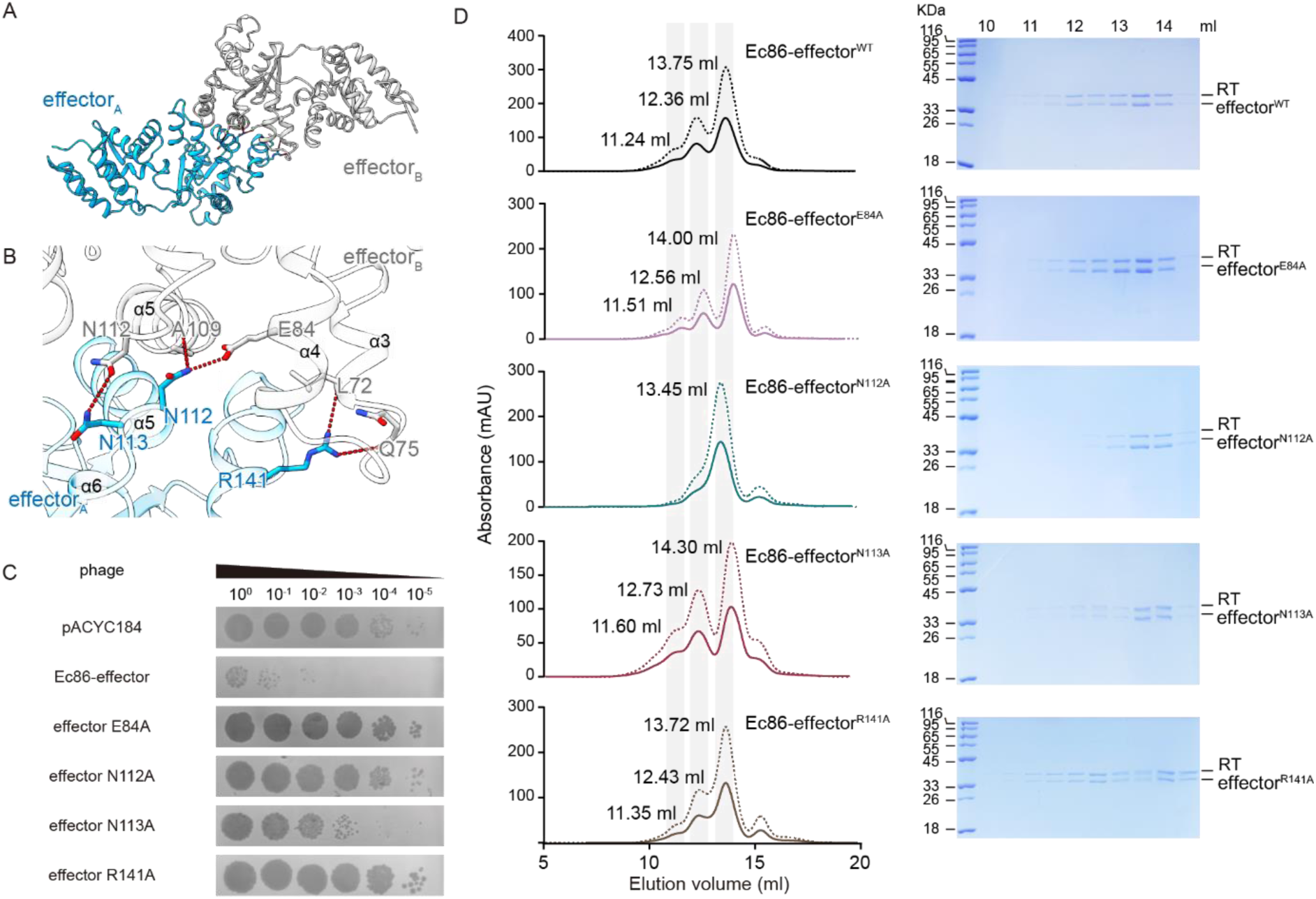
Effector-effector interactions in the retron filament are crucial for antiphage activity. (**A**) The effector-effector interaction interface in the Ec86-effector filament complex. The two effectors (effector_A_ and effector_B_) are shown in deep sky blue and gray, respectively. (**B**) A close-up view of the effector_A_-effector_B_ interaction interface. (**C**) Serial dilution plaque assays shown for the *E. coli* MG1655 strain transformed with plasmids encoding WT or mutated retron Ec86 systems. Images are representative of two replicates. (**D**) Purification of the WT or mutated Ec86-effector complex indicated by representative gel filtration chromatography. The absorbance at A280 (protein) and absorbance at A260 (nucleic acid) are indicated by solid line and dotted line, respectively. The elution peaks containing Ec86-effector complexes are illustrated by gray shadows. The elution fractions corresponding to the gel filtration chromatography are subjected to SDS-PAGE and Coomassie brilliant blue staining.

We then inspected the catalytic residue (E106) in both effectors and located two putative active-sites, designated active-site_A_ and active-site_B_, respectively, in effector_A_ and effector_B_ (Figure 5A). Notably, two putative ligands, with the cryo-EM densities assigned to a thymine and an adenosine triphosphate (ATP), respectively, were observed in the active-site_A_ that contains effector_A_ E106. In contrast, no additional densities were found in active-site_B_ (Figures 5A and 5B; Figure S7). For clarity, we named the pockets in the active-site_A_ accommodating the two putative ligands thymine-binding pocket and ATP-binding pocket, respectively. In the thymine-binding pocket, effector_A_ residues D69 and E106 tightly locks the thymine within the pocket, making it solvent-inaccessible (Figure 5C; Figures S7 and S8). The ATP-binding pocket is located at the asymmetric effector_A_-effector_B_ dimer interface, where a set of hydrophobic residues in effector_B_ were positioned to stack the adenine base of the ATP and a set of hydrophilic residues in both effector_A_ and effector_B_ were positioned to coordinate the phosphodiester linkage and ribose of the ATP. Specifically, effector_A_ residues E65, D69, and S100, as well as effector_B_ residues F128, K131, S133, and N136 were observed to interact with the ATP (Figure 5D). Further sequence and structure analysis of the Ec86 effector with the sequences and corresponding AlphaFold2 predicted structures of three other retron NDT-like effectors revealed that residues E65, S100, E106, and S133 are highly conserved (Figure S7). Alanine substitutions of these conserved residues that participate in substrate binding and/or catalysis resulted in notable deficiencies in antiphage activities (Figure 5E), confirming the critical roles of effectors’ ligands-binding residues. Taken together, these results indicate that the effector dimerization observed in the filament assembly is indispensable for Ec86-mediated antiphage defense.

**Figure 5.**
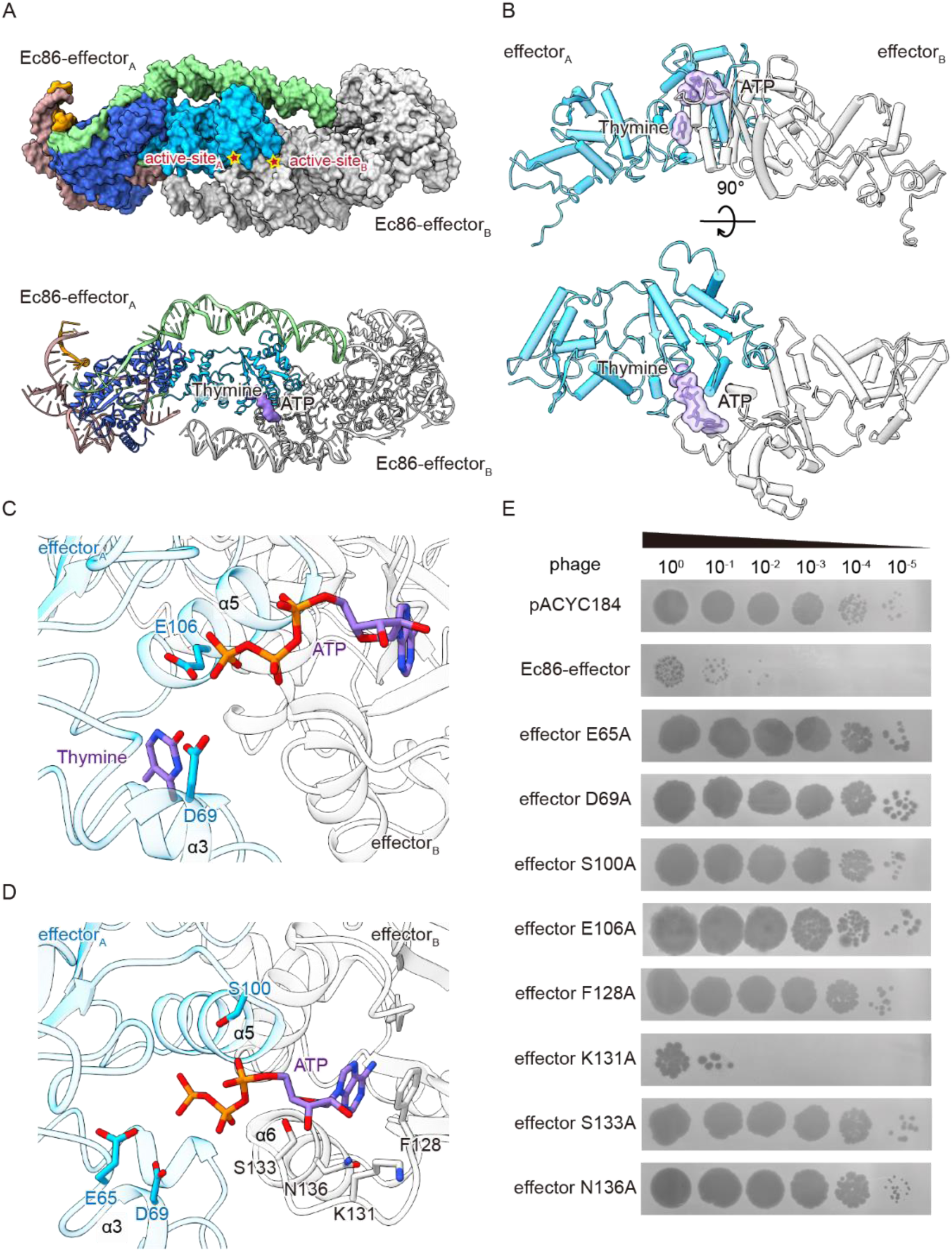
Effector active-site architecture in the Ec86-effector filament. (**A**) Surface (top panel) and cartoon (bottom panel) representations of two Ec86-effector subunits in the Ec86-effector filament complex. Red stars highlight the active-sites. The thymine and ATP shown in surface representation. (**B**) Cylinder representation of the effector dimer indicated in (A). The thymine and ATP shown in sticks and surface representation. (**C**) A close-up view of the interaction interface between the thymine and the effector dimer (deep sky blue and gray), with key residues shown as sticks. (**D**) A close-up view of the interaction interface between the ATP and the effector dimer (deep sky blue and gray), with key residues shown as sticks. (**E**) Serial dilution plaque assays shown for the *E. coli* MG1655 strain transformed with plasmids encoding WT or mutated Ec86 systems. Images are representative of two replicates.

### Activated Ec86 confers phage defense through depleting adenine and its derivatives

A defining feature of activation for classic deoxyribosyltransferases, such as the nucleoside 2-deoxyribosyltransferases from *Lactobacillus leichmannii* (*Ll*NDT) and *Bacillus psychrosaccharolyticus* (*Bp*NDT), is the formation of dimers or trimer-of-dimers^43,44^. Thus, we proposed previously that the Ec86 effector N-terminal NDT-like domain may form active dimers to exert its enzymatic activity for antiphage defense. In the Ec86-effector filament, the effectors indeed formed dimers; however, the toxicity (i.e., the activity of the effector) of the Ec86 system requires activation by phage factors. We thus hypothesize that the observed effector dimer may represent a state primed for activation rather than an active state. Structural comparison of the Ec86 effector dimer with an activated deoxyribosyltransferase (*Ll*NDT PDB: 1F8Y) revealed obvious differences in the active-site architecture (Figure S8A and S8B). Specifically, only one ligand can be found in *Ll*NDT active-site in-between α6 and α7 (containing catalytic residue E98), while two putative ligands can be observed in the Ec86 effector active-site between α3-loop-α4 and α5 (containing E106). The *Ll*NDT α6 is rigid and forms a pocket with α7 that holds the ligand, which is solvent-accessible (Figure S8C). In contrast, the Ec86 effector α3-loop-α4 region is more flexible than its *Ll*NDT counterpart (*Ll*NDT α6), and the space between effector α3-loop-α4 and α5 is bigger than that of the *Ll*NDT ligand-binding pocket. Besides, the ATP is exposed and solvent-accessible, whereas the thymine is buried within the active-site, locked by D69 and E106, and solvent-inaccessible (Figure S8D). These suggest that the Ec86-effector filament is not in an active state.

As the effector mechanism of the Ec86 system remains unknown, we further investigated the possible NDT-like enzymatic function of the Ec86 effector after activation by phage Dcm. We transferred plasmids that contain inducible Dcm or Dcm^mut^ into *E. coli* cells harboring the WT Ec86 system and collected cell lysates from several time points after induction. The cell lysates were then filtered and subjected to mass spectrometry analysis. We tested a panel of bases and nucleosides and observed that by 15 min after induction of Dcm expression, cell lysates contained significantly less amount of adenine, adenosine, and deoxyadenosine compared to the Dcm^mut^ and empty vector control samples (Figure 6A; Figure S9). This suggests that adenine and its derivatives are substrates of the Ec86 effector. Interestingly, we also observed the accumulation of guanine, guanosine, and deoxyguanosine after induction of Dcm expression (Figure 6B; Figure S9). No significant changes in cytosine, thymine, uracil, and their derivatives were observed (Figure S10). These results highlight that the Ec86-effector filament is primed for activation by phage Dcm and suggest that the activated Ec86 system mediates massive depletion of adenine (and adenine derivatives) and production of guanine (and guanine derivatives).

**Figure 6.**
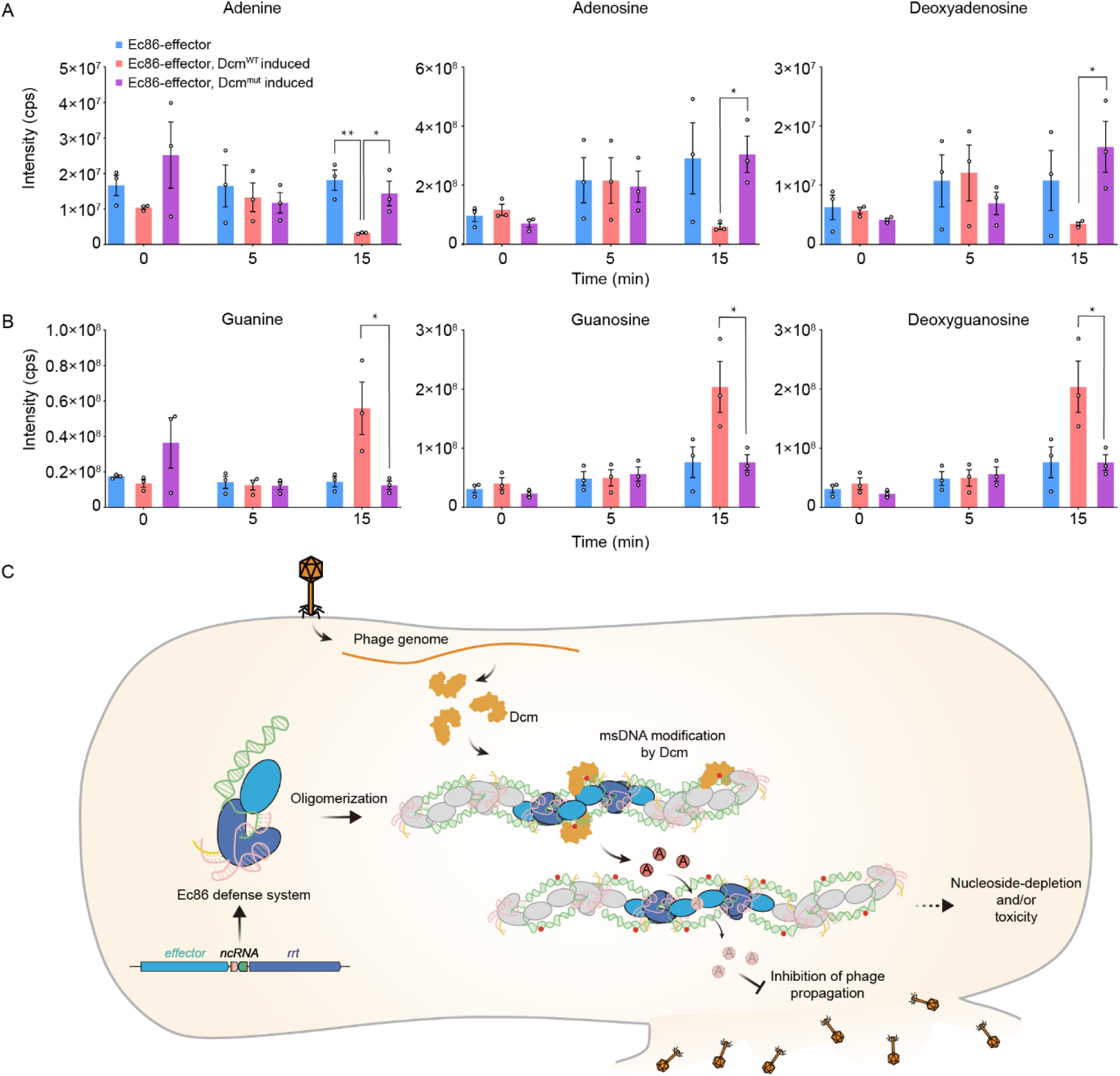
Activated retron Ec86 confers phage defense through nucleoside depletion. (**A-B**) In vivo abundance of adenine and adenine derivatives (**A**) and guanine and guanine derivatives (**B**) in *E. coli* MG1655 cells expressing the retron Ec86 system from its native promoter after induction of Dcm expression, Dcm^mut^ (that contains C82A, E128A, K175A, and K176A mutations) expression, or a corresponding empty vector. (Data represent the mean ± SEM of three biological replicate cultures. Unpaired t test. *, p < 0.05; **, p < 0.01) (**C**) Model for antiphage defense mediated by retron Ec86 system in bacteria. During phage infection, the phage-encoded Dcm methylates the msDNA DSL, leading to effector activation. The effectors then rapidly consume adenine derivatives, interfering with host nucleoside/nucleotide homeostasis and/or poisoning the nucleotide pool, which eventually results in the inhibition of phage replication and abortive infection.

## Discussion

In this study, we attempted to unveil the full picture of retron Ec86 system-mediated antiphage defense. Our results revealed that the retron defense system forms a supramolecular Ec86-effector filament assembly with effector dimers sandwiched by RT-msDNA. The filament assembly is primed for activation by phage Dcm-mediated msDNA methylation and confers defense through depletion of adenine and adenine derivatives upon activation. Considering that the DSL containing the Dcm methylation site directly contacts its cognate effector’s NDT-like domain, we thus propose the following model wherein the msDNA DSL in the Ec86 RT-msDNA functions as a phage sensor that activates the effector. Without phage infection, the NDT-like domains of DSL-bound effector dimers adopt an inactive conformation with closed active-site(s). During phage infection, phage-encoded Dcm methylates the msDNA DSL, affecting the DSL-effector interaction and presumably leading to the simultaneous conformational changes of the active-site(s) within the sandwiched effector dimer and effector activation. The effectors then rapidly consume adenine derivatives, interfering with host nucleoside/nucleotide homeostasis and/or poisoning the nucleotide pool, which eventually results in the inhibition of phage replication and abortive infection (Figure 6B).

Our results together with previous studies indicate that retron systems likely share a conserved phage recognition mechanism by which the DNA part of msDNA serves as the sensor of phage infection. Various triggers for retron defense systems have been identified recently^11,17,18,33,45^. Notably, almost all these identified triggers, including phage genes encoding DNA methylases, exonucleases, and single-stranded DNA-binding proteins, have the capacity or predicted capacity to directly interact with msDNA^17,18,33,45^. The only exception that does not seem to directly interact with msDNA is the phage-encoded inhibitors of RecBCD^11,17^. Possibilities are that the inhibition of RecB triggers the expression of another host DNA-binding protein that substitutes the role of RecB, which might directly interact with retron msDNA. In addition, another interesting finding is that both Dcm identified in this study and N6-adenine methyltransferase (Dam) identified previously are used by phages to defend against bacterial RM defense systems^2,33^. Phage-encoded RecBCD inhibitors circumvent the defense by the RecBCD system^46–49^. These suggest that retrons may sense immune evasion of early anti-phage defenses, including RM and RecBCD, and function as the second line of immunity^11,17,33^.

We identified the cellular target of the Ec86 effector and suggested that the activated system functions through the depletion of adenine derivatives. This marks an important effector mechanism via depletion of molecules essential for phage replication. Although not all effectors in the highly prevalent NDT-like-domain-containing retron family exhibit obvious toxicity^33,50^, it is possible that they function in a manner similar to the Ec86 effector after activation. More broadly, various antiviral defense systems, including prokaryotic RADAR, AvcID, and eukaryotic SAMHD1, function through nucleotide depletion^25–28,51,52^, highlighting a biologically conserved antiviral strategy of targeting the cellular nucleotide pool and limiting virus reproduction^25,26,28^. Apart from this, Ec86 effector NDT-like domains couple the scaffolding function that mediates the formation of protein filaments (supramolecular complex assembly) with the enzymatic activity required for antiphage defense. Extensive multimerization of repeating subunits in Ec86 defense may specifically enhance the effector’s enzymatic activity required for anti-phage defense. The Ec86-effector filament, together with prokaryotic supramolecular complexes (in CRISPR, CBASS, RADAR, Gabija, AVAST immunity), mammalian inflammasomes, and plant resistosomes, highlights an emerging theme of supramolecular complex assemblies as a critical step in innate immunity^10,15,27,28,53–62^.

## Acknowledgments

We thank the cryo-EM Facility of University of Science and Technology of China for providing cryo-EM and computation support and the Center for Protein Research and Public Laboratory of Electron Microscopy, Huazhong Agricultural University, for technical support; Yongxiang Gao, and Jianbo Cao for help with cryo-EM data collection; Erchao Sun for technical support in cloning and phage plaque assays; Hongbo Liu and Junjun Yan for technical assistance, data collection, and data analysis with the mass spectrometry experiments.

## Author contributions

T.Z. conceived the project. Y.W., C.W., P.T., and T.Z. designed all experiments. Y.W., C.W., J.C., J.X., S.W., Y.Cui., Z.Q., and Y.Chen. performed the experiments. Q.W. prepared the cryo-EM samples and collected cryo-EM data. Z.G. determined the structure. All authors analysed the data and contributed to manuscript preparation. Y.W., C.W., and T.Z. wrote the manuscript.

## RESOURCE AVAILABILITY

### Materials availability

Bacterial and virus strains generated in this study are available from the lead contact upon request.

### Data and code availability

- Raw sequencing data have been deposited to NCBI which is publicly available as of the date of publication.
- This paper does not report original code.
- Any additional information required to reanalyze the data reported in this paper is available from the lead contact upon request.

## EXPERIMENTAL MODEL AND SUBJECT DETAILS

### Bacterial strains and phages

*Escherichia coli* strains [MG1655, DH5α, and BL21(DE3)] were grown in LB or LB agar at 37°C shaking at 200 rpm unless mentioned otherwise. Whenever applicable, media were supplemented with ampicillin (100 μg ml^-1^), kanamycin (50 μg ml^-1^) or chloromycetin (25 μg ml^-1^) to ensure the maintenance of plasmids. The Hzau phage collections were isolated from sewage samples on *E. coli* MG1655. Infection was performed in LB medium supplemented with 0.1 mM MnCl_2_ and 5 mM MgCl_2_ at 37 ℃.

### Method details

#### Plasmid construction

The full-length Ec86/effector genetic unit (*effector-ncRNA-rrt*) gene with its native promoter was amplified from its native host *E. coli* BL21 (DE3) and subcloned into pACYC184 vectors for phage challenge assays. The Ec86/effector genetic unit (*effector-ncRNA-rrt*) with a C-terminal 8×His tag (His_8_) was introduced into pET21b vector (Novagen) for expression and purification. The mutants Ec86/effector genetic unit were generated using the original sequence as the template. The full-length *dcm* gene was cloned from phage K42 and subcloned into pBAD vector for in vivo experiments.

#### Isolation of phage K42 from sewage

Phage K42 was isolated from sewage as previously described^1,63^. Sewage samples were filtered concentrated with 0.2 μm filter, and subsequently used to perform double layer plaque assays. Individual phage plaques were picked, reisolated 3 times, and amplified as described below.

#### Isolation of mutant phages

To isolate mutant phages that escape Ec86 defense system, phages were plated on bacteria expressing the Ec86 defense system using the double-layer plaque assay^64,11^. MMB (LB + 0.1 mM MnCl_2_ + 5 mM MgCl_2_) 1.1% agar and 0.5% agar were used as the bottom layer and the top layer, respectively. For this, 400 ml *E. coli* MG1655 containing Ec86 defense system grown in LB medium until optical density at 600 nm (OD_600_) reached 0.3, and then, bacteria mixed with 7ml pre-melted 0.5% MMB agar poured onto MMB 1.1% agar plates. After drying 10 minutes at room temperature, 2 ml drops of the tenfold serial diluted phage k42 culture was spotted on the bacterial layer. The double layer plates were incubated overnight at 37 ℃, and single plaques were picked into 90 ml phage buffer. The phages were mixed several times by vortex to release them from the agar into the phage buffer, after which the phages were centrifuged at 2000 g for 10 min to get rid of agar and bacterial cells, and the supernatant was transferred to a new tube. To test whether the phages can escape from Ec86 defense, the small drop plaque assay was used as described below.

#### Plaque assays

Plaque assays were performed as previously described^64,11,3^. In brief, *E. coli* MG1655 transformed with pACYC184 plasmids (containing different constructs) or empty vector were grown overnight at 37°C with shaking at 210 rpm. After overnight culture, 400 μl bacteria was mixed with 7 ml molten top agar. The mixture was poured onto LB agar plates. Phage solution (wild type phage or mutant phages) was subjected to 10-fold serial dilutions in LB medium. After the top agar dried, 2 μl drops of the diluted phage culture were spotted on the top bacterial layer. The plates were imaged after overnight incubation at 37°C. Efficiency of plating (EOP) was measured and compared between the defense and control strain.

#### Amplification of phages

Amplification of phages were performed as previously described^11,64^. *E. coli* MG1655 was cultured in MMB medium at 37°C with shaking at 210 rpm overnight. Then bacteria were diluted 1:50 in MMB medium and grown to OD600 0.3. The phage k42 were propagated by mixing 10 μl phage with 500 μl of the bacterial culture, and incubated for 3 h at 37 ℃ with shaking at 200 rpm. The lysate was centrifuged for 10 min at 2,000 ×g and the supernatant were filtered using a 0.22 μM filter to get rid of remaining bacterial cell debris. The phage titer was then determined using the small drop plaque assay. Wild type phage or mutant phages were propagated in *E. coli* MG1655 with empty vector or defense-system-expressing *E. coli* MG1655, respectively.

#### Sequencing and genome analysis of phage mutants

The extraction of phage genome were performed as previously described^11,16,17^. High titer phage lysates (> 10^7^ pfu ml^-1^) of the ancestor and isolated phage mutants were used for DNA extraction. 5 ml of the phage lysate was treated with 20 μl DNase-I (1 mg ml^-1^) and 5 μl RNase-A (10 mg ml^-1^) at 37°C for 1 hour to remove bacterial DNA. Subsequently, 20 μl 2 M ZnCl_2_ was added and incubated at 37°C for 5 min. Samples were centrifuged at 12,000 rpm for 1 min. Pellets were resuspended by 500 μl TES buffer (0.1 M Tris-HCl pH 8.0, 0.1 M EDTA and 0.3% SDS) at 65°C for 15 min. Lysates were then incubated with proteinase K at 50°C for 1.5 hours to disrupt capsids and release phage DNA, and subsequently added 60 μl pre-cold CH3COOH (3 M, pH 5.2) on ice for 15 min. Samples were centrifuged at 13,000 rpm for 10 min at 4°C. The supernatant was extracted with 600 μl phenol:chloroform:isoamyl alcohol (25:24:1), and phage DNA was precipitated overnight at 4°C with isopropanol at -20°C. The precipitated phage DNA was centrifuged at 12,000 ×g for 1 hr at 4°C and the pellet was washed twice with 1 ml 70% ethanol. The pellets were air-dried for 15-30 min at room temperature and resuspended in 30 μl 1×TE buffer (10 mM Tris-HCl, 0.1 mM EDTA, pH 8.0). Concentrations of extracted DNA were measured by NanoDrop (Thermo Fisher Scientific). Library preparation and whole-genome sequencing using Illumina HiSeq sequencing platform with a PE150 cartridge was performed by the Annoroad Biotechnology Co., Ltd (Zhejiang). Quality trimming was performed using Trimmomatic v. 0.39 (LEADING:3 TRAILING:3 SLIDINGWINDOW:4:15 MINLEN:36)^65^. Trimmed reads were assembled de novo using SPAdes v. 3.13.0 with default parameters and k-mer lengths of 21, 33, 55 and 77^ref66^. The assembled genomes were polished using Pilon v1.2.4^ref67^. Classification of the phage family was performed according to the most closely related known phage based on sequence similarity, as described previously^1^. Only mutations that occurred in the isolated mutants, but not in the ancestor phage, were considered. Silent mutations within protein coding regions were disregarded as well. Raw sequencing data was deposited to NCBI.

#### Bacterial growth upon induction of *dcm*

*E. coli* Mg1655 were co-transformed with pACYC184 vector (containing Ec86 operon or empty vector) and arabinose induced pBAD vector (containing wild type or mutant *dcm*), and grown in LB (supplemented with ampicillin and chloromycetin) at 37 ℃ with shaking at 200 rpm overnight. Bacteria were diluted 1:50 into 5 ml LB media. 180 μl of the culture were dispensed into a 96-well plate and incubated at 37 ℃ with shaking in a Microplate Reader (FLUOstar® Omega) until OD_600_ reached 0.3. 20 ml of ultra-pure water (for uninduced samples) or 2% arabinose (for induced samples, for a final concentration of 0.2%) was added to the bacterial cultures and the bacteria were then incubated further at 37 ℃ with shaking in a Microplate Reader (FLUOstar® Omega) with OD600 measurement every 6 minutes.

#### Spot growth tests

Spots in growth tests performed as previously described^33^. For *E. coli* MG1655, single bacterial colonies containing pACYC184 vector (containing Ec86 operon or empty vector) and arabinose induced pBAD vector (containing wild type or mutant *dcm*), were picked into LB supplemented with ampicillin and chloromycetin, and cultured at 37℃ with shaking at 200 rpm. When bacteria reached an OD600 of 0.8, cultures were stepwise serially diluted 8 times (tenfold) in LB (100 μl culture + 900 μl LB). 2 μl of culture dilutions were spotted on LB plates containing ampicillin, chloromycetin and 0.2% arabinose (for final concentration) and incubated at 37℃ overnight.

For *E. coli* BL21 (DE3, the native strain of Ec86 defense system), bacteria were transformed with pBAD vector (containing wild type or mutant *dcm*), and grown in LB at 37 ℃ with shaking at 200 rpm for 2 hrs. Cultures were 100-fold diluted and poured on the LB agar media with or without 0.2% arabinose (final concentration). LB plates were incubated overnight at 37°C. Colony-forming units (CFU) was measured.

#### Detection of methylation of msDNA upon induction of *phage genes*

Ec86 msDNA was isolated by alkaline lysis as previously described^11,33^. In brief, msDNA was produced by co-transferring pACYC184-*effector-ncRNA-rrt* and arabinose-induced pBAD plasmids (containing *dcm^WT^*, *dcm^mut^*or empty vector) into *E. coli* MG1655 strain, in order to detect whether msDNA was methylated by Dcm. Strains were inoculated in 25 ml LB supplemented with ampicillin and chloromycetin, and incubated for 1.5 h at 37°C with shaking at 200 rpm. When bacteria reached an OD600 of 0.3, cultures were induced by the addition of 0.2% arabinose, and incubated for a further 5 hrs. After this, samples were centrifuged for 10 minutes at 2,000 ×g at 4°C, and the pellet was retained.

To isolate the msDNA, each pellet was treated with 200 μl Solution I (50 mM glucose, 10 mM EDTA and 25 mM Tris-HCl, pH 8.0), 200 μl Solution II (0.2 N NaOH, 1% SDS) and 400 μl Solution III (3 M potassium acetate, 2 M acetic acid). The tubes were mixed by inverting each tube 5 times after each solution added. Samples were centrifuged for 10 minutes at 4000 rpm at 4°C, after which the supernatant was collected.

Nucleic acids were subsequent overnight precipitated with 3 volumes of 100% ethanol and 10% (v/v) 3M sodium acetate (NaAc) pH 5.2 at -80°C. The precipitated nucleic acids were centrifuged at 12,000 ×g for 1 hr at 4°C and the supernatant was discarded. The pellet was washed twice with 1 ml 70% ethanol. After centrifugation at 12,000 ×g for 10 min at 4°C, the supernatant was discarded, and the pellets were air-dried for 15-30 min at room temperature and resuspended in 20 μl DEPC-treated water. The purified nucleic acids containing msDNA were detected by denaturing urea polyacrylamide gels electrophoresis (urea-PAGE, 12%) with 1×TBE buffer (150 volts, 1 hour).

To detect the methylation status of msDNA, the extracted msDNA was further purified from RNA and DNA contaminants by the crush and soak method^52^. In brief, extracts (from 25 ml of bacterial cultures) were digested by 5 μl RNase-A (10 mg ml^-1^) for 1 h to remove RNA part of msDNA and RNA contaminants, and subsequently loaded per well in 8% TBE-polyacrylamide gels (150 V for 1 h). Gels were stained with 4S GelRed, and the msDNA-containing-gel slices were transferred to 1.5 ml tubes. The gel slices were crushed against the walls of the tubes with a tip, suspended in two gel-slice volumes of 0.5M ammonium acetate pH 5.2, and incubated at 37°C with shaking at 200 rpm for 2 h. Samples were centrifuged at 14,000 rpm for 10 min at room temperature, and the supernatants were transferred to fresh tubes. 3 volumes of 100% ethanol and 10% (v/v) 3M sodium acetate (NaAc, pH 5.2) was added to the supernatants, and msDNA was precipitated overnight at -80°C. Samples were centrifuged at 14,000 rpm for 60 min at 4°C, pellets were washed once with 1 ml of 70% ethanol, and washed pellets were centrifuged again at 14,000 rpm for 10 min at 4°C. Pellets (purified msDNA) were air-dried for 30 min, and resuspended in 12 μl of diethyl pyrocarbonate-treated water. The samples detected by 8% TBE-polyacrylamide gels and quantified by Nanodrop. To hydrolyse DNA part of msDNA into nucleoside, 0.5 μl P1 nuclease (1 mg ml^-1^, sigma) and 0.5 μl Calf Intestinal Alkaline Phosphatase (CIAP, 3 U μl^-1^, Invitrogen) was added in 15 μl of purified msDNA and digested 3 hrs at 37°C. The samples were centrifuged at 14,000 rpm for 10 min at 4°C. Filtrate was taken and used for LC-MS analysis.

Experimental samples and standards 5mC (macklin) analysis was performed on a triple Quad 5500 (AB SCIEX, USA) LC-MS/MS system. A ODS C-18 (Shimadzu Corporation) column (2.1 × 150 cm, 3 μm) was used at a flow rate of 0.25 ml min^-1^. The injected volume of sample was 10 μl. The elution gradient was carried out with binary solvent system consisting of water (solvent A) and acetonitrile (solvent B). A linear gradient profile with the following proportions (v/v) of solvent B was applied: gradient profile 0 to 0.5 min and 5% of B, 0.5 to 10 min and 5% to 50% of B, 10 to 12 min and 50% of B, 12 to 13 min and 50% to 95% of B, 13 to 15min and 95% of B, 15 to 15.1 min and 95% to 5% of B, and 15.1 to 20 min with 5 min for re-equilibration and 5% of B. The MS spectra were processed with Analyst TF 1.6.2 (SCIEX, USA). MS conditions were as follows: source voltage 4.5 kV, source temperature 550, gas 1 and 2, nitrogen 55 psi and 50 psi; curtain gas, nitrogen 35 psi.

#### Protein expression and purification

Protein expression and purification were performed as previously described^34^. The pET21b-*effector*-*ncRNA*-*rrt*-His_8_ was transformed into *E. coli* BL21(DE3) Δ*Ec86/effector* genetic unit (i.e., Δ*effector*-*ncRNA-rrt*) strain for the expression of the Ec86-effector complex. Pre-cultures inoculated with a single colony were grown in LB (Luria-Bertani) medium supplemented with 100 μg ml^-1^ ampicillin at 37°C with shaking overnight. The pre-culture (10 ml) was added into 1 L LB medium (supplemented with 100 μg ml^-1^ ampicillin) and incubated at 37°C with shaking at 200 rpm until optical density at 600 nm (OD_600_) reached 1.0. At this point, the temperature was lowered to 16°C. The cultures were induced by the addition of 200 μM isopropyl-β-D-thiogalactoside (IPTG) and incubated for a further 16 hrs. Then cells were harvested by centrifugation.

For protein purification, the collected cells were resuspended in Lysis Buffer (25 mM Tris-HCl, pH 8.0, 500 mM NaCl) and subsequently disrupted by pressure homogenizer at 4°C. The lysate was clarified by centrifugation at 20,000 ×g for 1 hr at 4°C. The supernatant was loaded onto a Ni^2+^ affinity resin (Ni-NTA, Qiagen), and eluted with buffer E (25 mM Tris-HCl, pH 8.0, 250 mM imidazole, 500mM NaCl). The eluted sample was applied to a Source15Q column (GE Healthcare), followed by a gradient NaCl elution from 500 mM to 1 M NaCl in 25 mM Tris-HCl, pH 8.0 and 2 mM 1,4-dithiothreitol (DTT). The fractions of elution peak were concentrated and subsequently purified by gel filtration chromatography (Superose6 10/300, GE Healthcare) in buffer SD (25 mM Tris-HCl, pH 8.0, 350 mM NaCl, 5 mM DTT). The UV absorbance values at 260 nm and 280 nm were detected. The purity of the effector-bound Ec86 complex was checked by SDS-polyacrylamide gel electrophoresis (SDS-PAGE).

#### Cryo-EM grid preparation and data acquisition

Aliquots (4.0 µl) of the freshly purified Ec86-effector filament (∼0.5 mg ml^-1^) were dropped onto glow discharged holey carbon grids (Quantifoil Au R1.2/1.3, 300 mesh). After blotting for 4.5 s with 100% humidity at 8°C, the grids were plunge-frozen in liquid ethane cooled by liquid nitrogen in Vitrobot (Mark IV, Thermo Fisher Scientific). The cryo-grids were transferred to 300 kV Titan Krios electron microscopes (Thermo Fisher) equipped with a GIF Quantum energy filter (slit width 20 eV) and a Gatan K3 Summit detector. Micrographs were recorded in the counting mode with a magnification of 81,000 ×, resulting in a pixel size of 1.07 Å. Each micrograph stack, which contains 32 frames, was exposed for 3.5 s with a total electron dose of 55 e^-^ Å^-2^. EPU software (version 2.9) was used for fully automated data collection. All images were motion-corrected using MotionCor2 with a binning factor of 1 with dose weighting. The defocus value of each image was set to -1.2 ∼ -2.2 μm and determined by Gctf.

#### Cryo-EM data processing

A diagram of the procedures for data processing is presented in Figure S3. For the oligomeric effector-bound Ec86 complex, 3,098 movies were collected and motion-corrected using MotionCor2. 3,098 well quality micrographs were selected, from which a total of 3,107,964 particles were picked automatically picked using the cryoSPARC v4.0 Blob picker^68^ and relion v4.0^69^ for 2D and 3D classification. After several rounds of 3D classification, 270,375 particles belonging to the best class were selected and subjected to 3D refinement. A cryo-EM density of the oligomeric effector-bound Ec86 complex with an estimated resolution of 2.7 Å based on gold-standard Fourier shell correlation (FSC)^70^ was obtained after non-uniform refinement and local refinement. DeepEMhancer was applied to modify the obtained cryo-EM map^71^. Local resolution variations of above cryo-EM map were estimated using Reamap^72^.

#### Model building and refinement

For atomic model of the oligomeric effector-bound Ec86, the optimized model effector-bound Ec86 complex described previously was rigid docked into the cryo-EM density in Chimera^34^. The initial model was further adjusted manually using COOT v0.8.9.1^73^. The resulting protein model was subsequently used for the rigid fit into the density of the entire cryo-EM density map via the phenix.dock_in_map function of the PHENIX v1.13 software suite^74^. Model quality was evaluated using the Molprobity scores^75^, the Ramachandran plots and EMRinger^76^. Figures were generated using ChimeraX (version 1.2.5) and PyMol (version 2.5.1).

#### Cell lysate preparation for trigger induced

Overnight cultures of *E. coli* co-transferring pACYC184-*effector-ncRNA-rrt* and arabinose-induced pBAD plasmids (containing *dcm^WT^*, *dcm^mut^*or empty vector) were diluted 1:50 in 200 ml LB medium and grown at 37°C (200 rpm.) until reaching OD600 of 0.3. The cultures were induced by 0.2% arabinose. Following the addition of arabinose, at 0, 5, and 15 min post-induction (plus an uninducible control sample), 30 ml samples were taken and centrifuged for 10 min at 2,000g. Pellets were flash frozen using liquid nitrogen. The pellets were re-suspended in 400 μl of 350 mmol l^−1^ ammonium acetate buffer at pH = 7 and supplemented with 4mg ml^−1^ lysozyme. The samples were then lysed using a tissue-lyser (Shanghai Jingxin industrial development C0., Ltd.) for 90 s at 60 Hz (three cycles). Tubes were then centrifuged at 4°C for 10 min at 14,000g. Supernatant was transferred to Amicon Ultra-0.5 Centrifugal Filter Unit 3 kDa (Merck Millipore, catalogue no. UFC500396) and centrifuged for 45 min at 4°C at 12,000g. Filtrate was taken and used for LC-MS analysis.

#### Quantification of nucleotides by HPLC-MS

Quantification of nucleotides was analysed by HPLC-MS using a triple Quad 5500 (AB SCIEX, USA) LC-MS/MS system. A ODS C-18 (Shimadzu Corporation) column (2.1 × 150 cm, 3 μm) was used at a flow rate of 0.25 ml min^-1^. The injected volume of sample was 10 μl. The elution gradient was carried out with binary solvent system consisting of water (solvent A) and acetonitrile (solvent B). A linear gradient profile with the following proportions (v/v) of solvent B was applied: gradient profile 0 to 2 min and 5% of B, 2 to 8 min and 5% to 16% of B, 8 to 9 min and 16% of B, 9 to 12 min and 16% to 90% of B, 12 to 15min and 90% of B, 15 to 15.1 min and 90% to 5% of B, and 15.1 to 20 min with 5 min for re-equilibration and 5% of B. The MS spectra were processed with Analyst TF 1.6.2 (SCIEX, USA). MS conditions were as follows: source voltage 5.5 kV, source temperature 550, gas 1 and 2, nitrogen 55 psi and 60 psi; curtain gas, nitrogen 30 psi. Adenine, adenosine, deoxyadenosine, guanine, guanosine, deoxyguanosine, cytosine, cytidine, deoxycytidine, thymine, deoxythymidine, uracil and uridine were purchased from macklin used as deoxy-nucleosides, nucleosides, and bases standards.

#### Quantification and statistical analysis

Details for specific statistical tests are found in the legends associated with each figure. In brief, data represent an average of three biological replicates, each with three technical replicates unless stated otherwise. Bars indicate mean ± SEM. The unpaired t test was conducted using Graphpad Prism v. 8.0.2 software to determine significant differences between groups. The p values represented in each figure are shown in the figure legends.

---

**Fig. S1.**
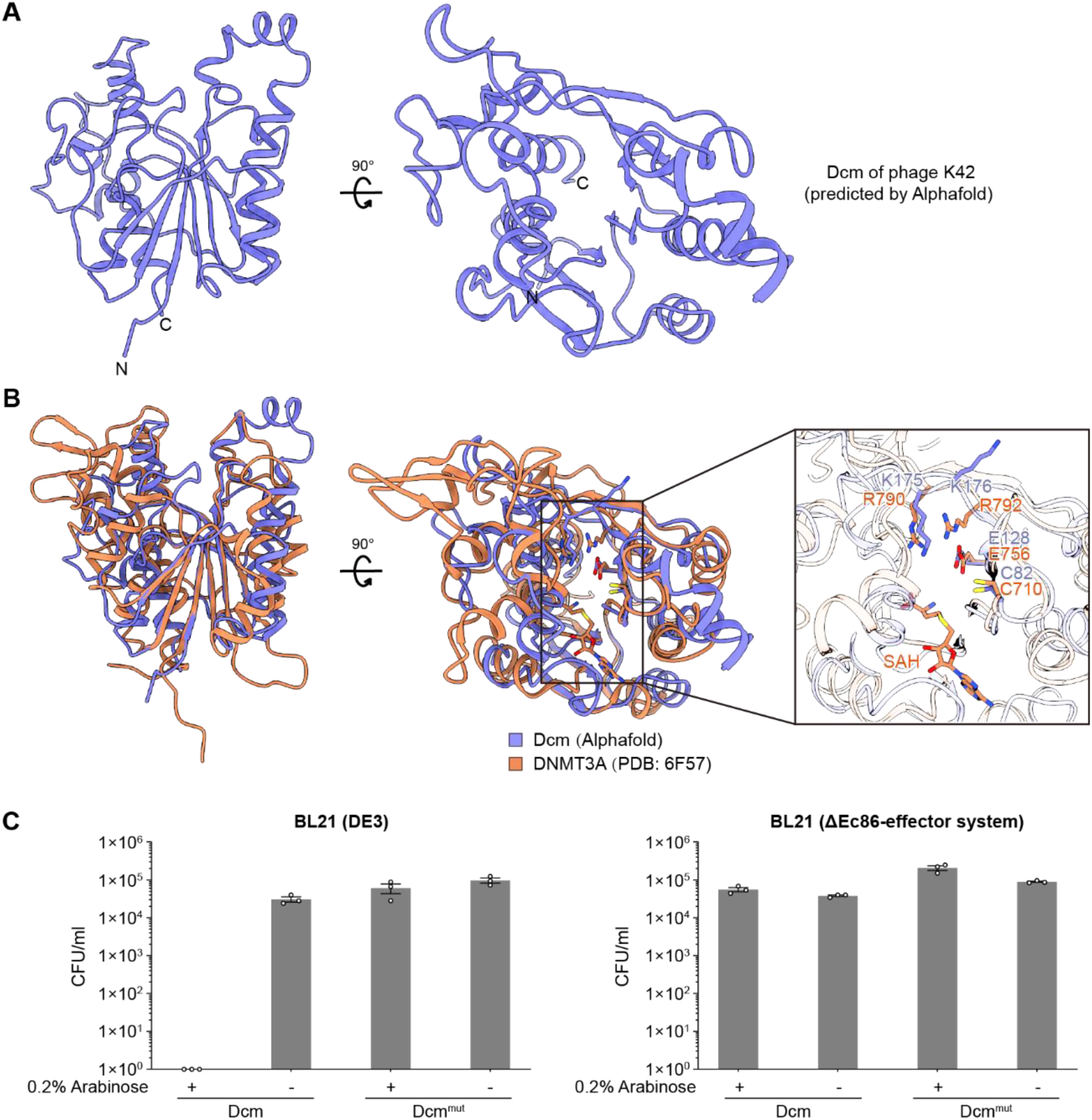
Superposition of the phage-encoded Dcm with DNMT3A. (**A**) AlphaFold2 predicted structure of phage K42-encoded Dcm. (**B**) Superposition of the predicted Dcm structure with DNMT3A (PDB: 6F57; RMSD= 4.364369 Å). Right panel, the close-up view of the DNMT3A active site and the predicted Dcm active site. The putative Dcm catalytic residue (cysteine 82, C82) and substrates binding residues (E128, K175 and K176) in the active site are depicted as sticks. DNMT3A, coral; Dcm, medium slate blue.(**C**) Overexpression of Dcm in *E.coli* BL21(DE3) that contains a native retron Ec86 operon inhibits bacterial growth. *E. coli* BL21(DE3) containing Dcm^WT^ or Dcm^mut^ (that contains C82A, E128A, K175A, and K176A mutations) were grown for 4 h in LB, serially diluted, spotted on LB plates containing no or 0.2% arabinose, and incubated at the 37℃. The number of colony forming units following transformation is presented. Bar graph represents an average of three biological replicates, with individual data points overlaid. Data represent the mean ± SEM of three biological replicate culture.

**Fig. S2.**
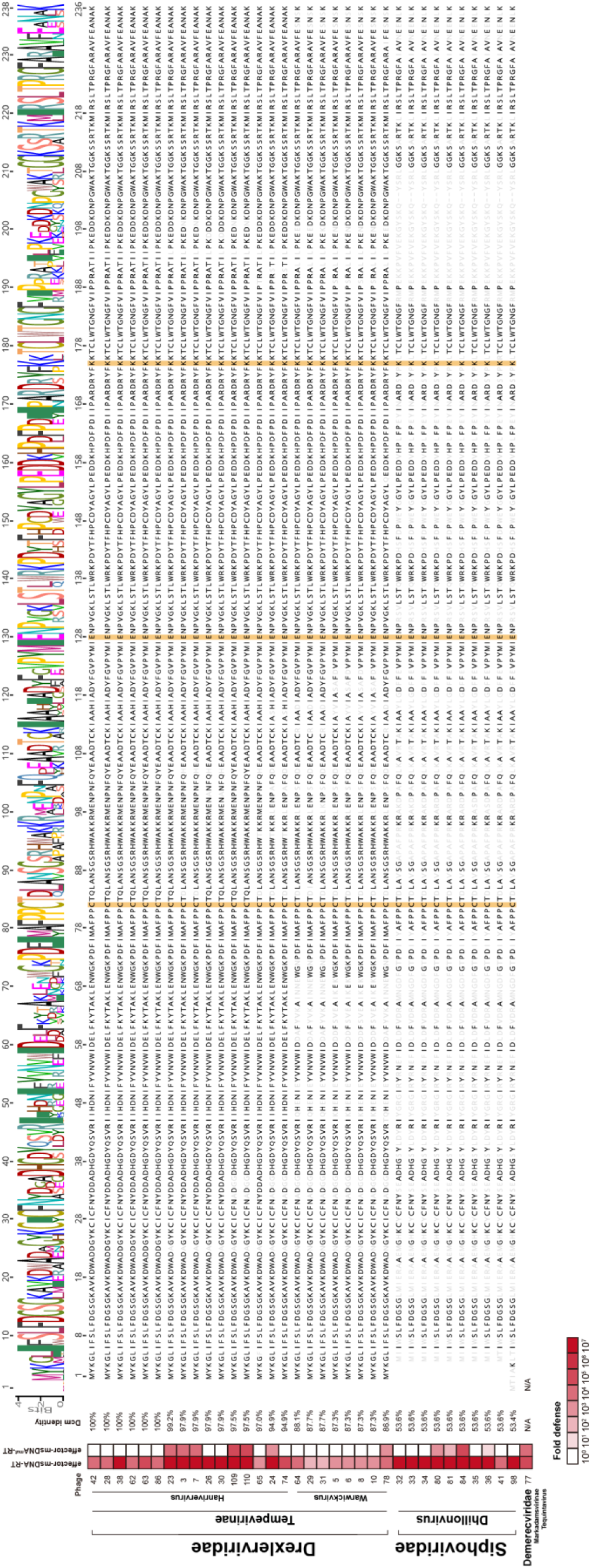
Overview of the retron Ec86-sensitive Hzau phage collection, their Dcm amino acid sequences, and Dcm identity compared to phage K42. Fold protection was measured using serial dilution plaque assays, comparing the efficiency of plating (EOP) of phages on the system-containing strain to the EOP on a control strain that lacks the system. Data represent an average of three replicates. The mutated Ec86 system contains msDNA^mut^ with the 5′-AATGG-3′ motif instead of the 5′-CCTGG-3′ motif. The conserved Dcm SAM-binding and catalytic residues are indicated by orange shadows.

**Fig. S3.**
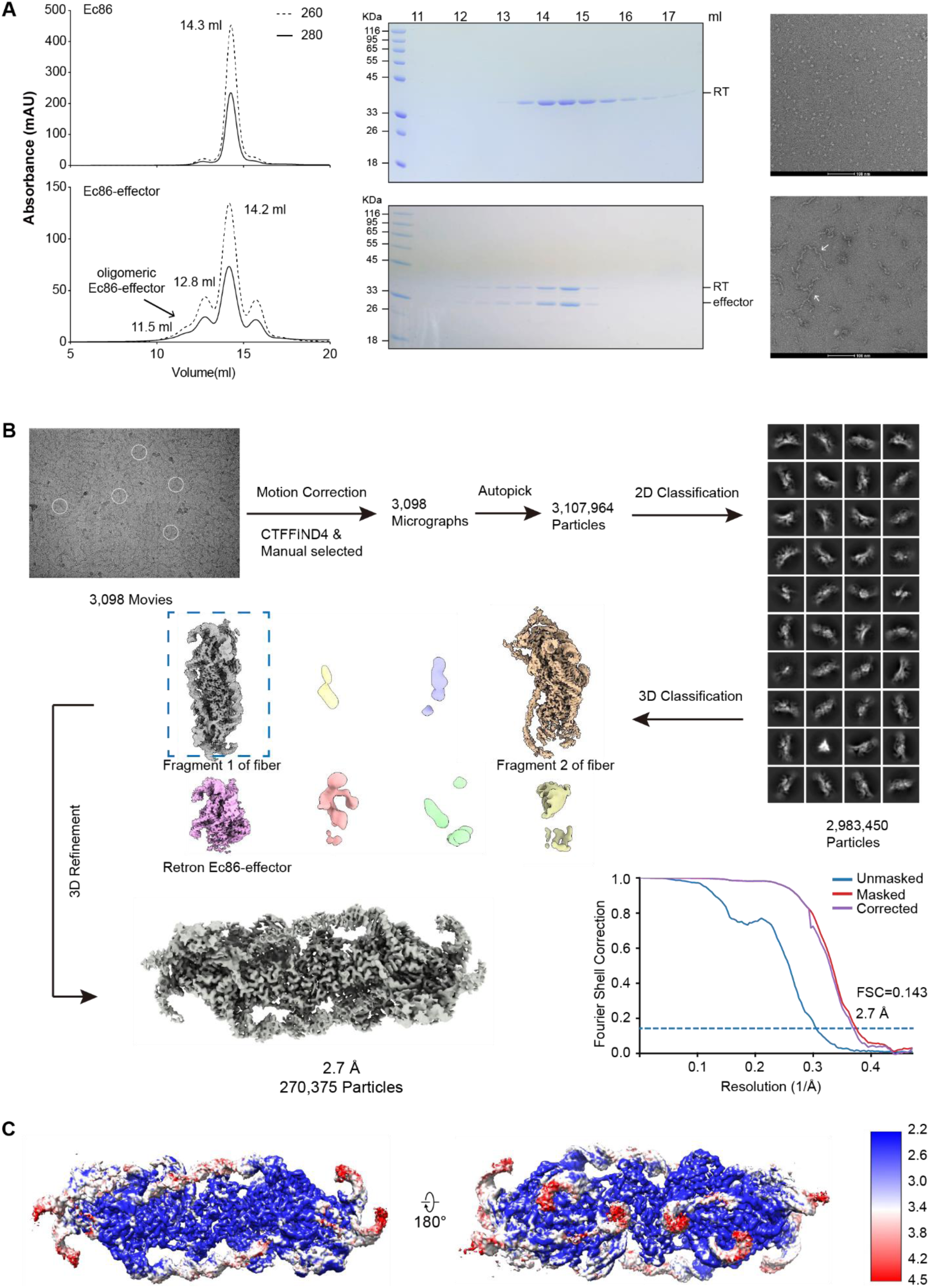
Purification and cryo-EM single particle analysis of the Ec86-effector filament. (**A**) Left, purification of the Ec86-effector filament is indicated by a representative gel filtration chromatography and SDS-PAGE. The peaks containing the target complex are illustrated by a black arrow. Images are representatives of three independent experiments. Right, Visualizations of the samples from the peak fractions using negative-stain EM. (**B**) Flowchart for the cryo-EM data processing. See Methods for detailed information. (**C**) Local map resolutions of the final structure.

**Fig. S4.**
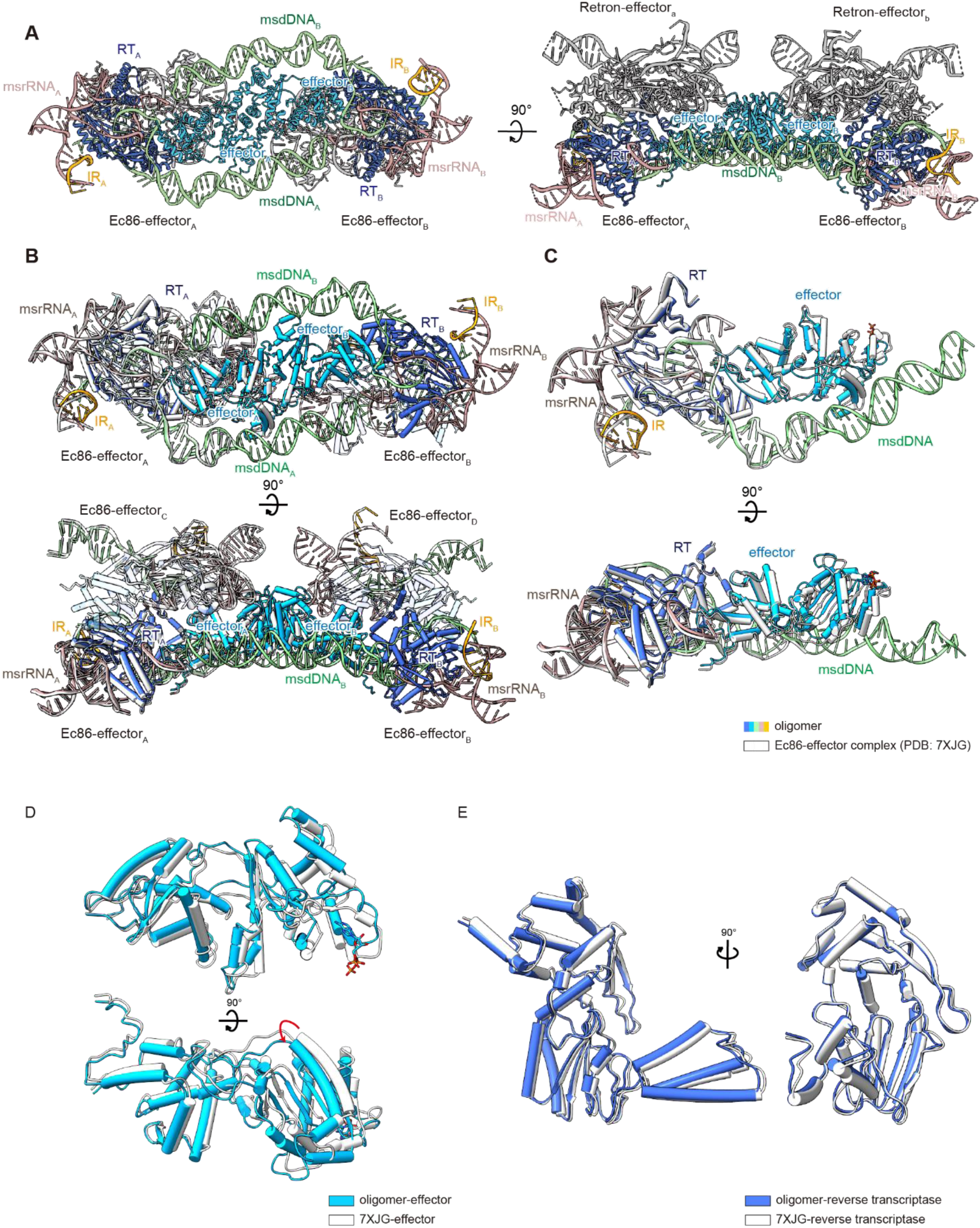
Comparison of the Ec86-effector filament and the dimeric effector-bound Ec86. (**A**) Cartoon representation of the Ec86-effector filament complex. (**B**) and (**C**) Superposition of the dimeric effector-bound Ec86 (PDB: 7XJG) and a repeating unit of the Ec86-effector filament structures (RMSD=1.19). (**C**) Superposition of the individual effectors from effector-bound Ec86 (PDB: 7XJG) and a repeating unit of the Ec86-effector filament structures. (**D**) Superposition of the RTs from effector-bound Ec86 (PDB: 7XJG) and a repeating unit of the Ec86-effector filament structures.

**Fig. S5.**
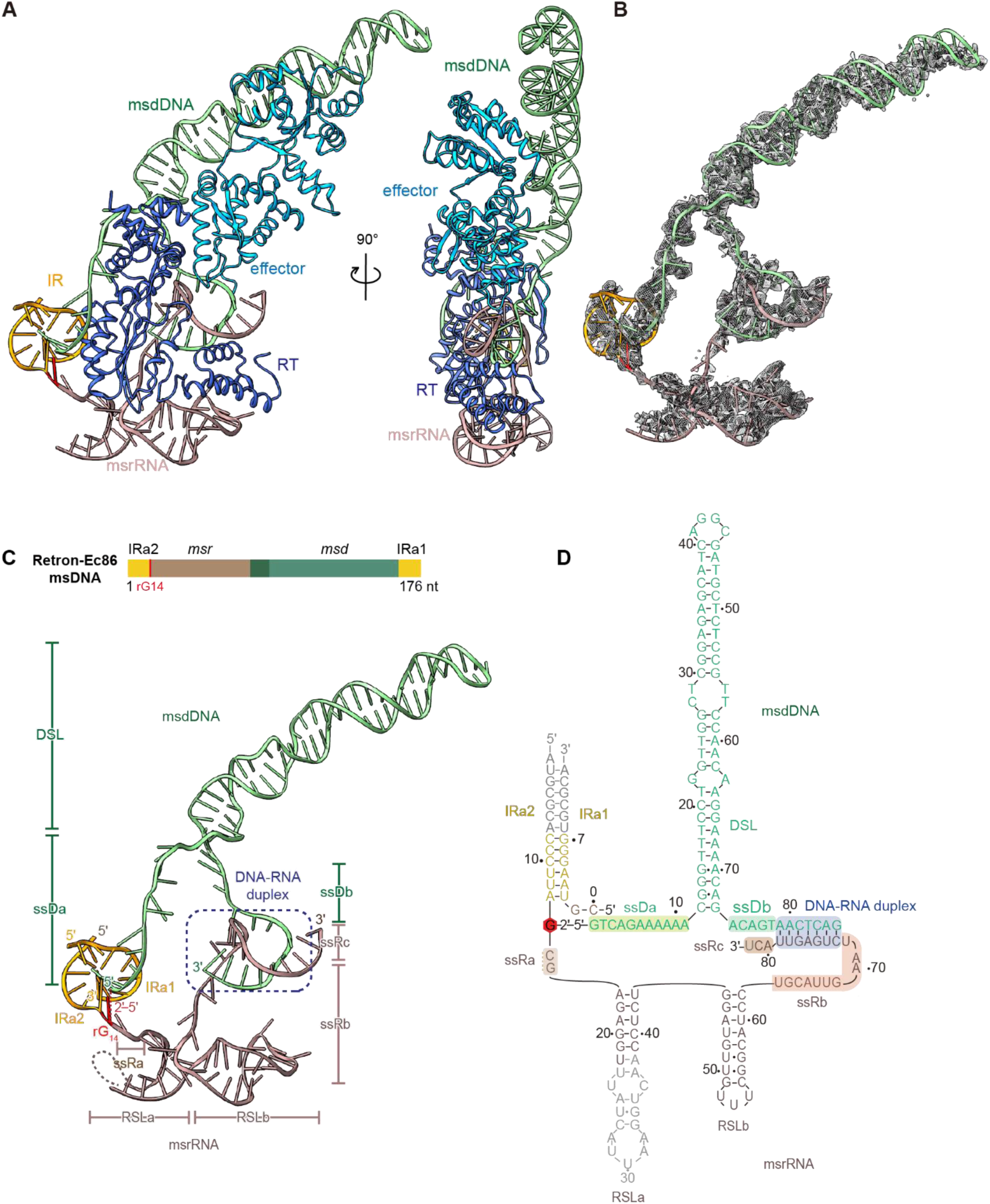
Structure of the intact Ec86 msDNA in a repeating unit of the Ec86-effector filament. (**A**) Atomic model of the Ec86-effector protomer from a repeating unit of the Ec86-effector filament in cartoon. (**B**) Cryo-EM map of the intact msDNA. (**C**) Top, schematic representation of the Ec86 msDNA. Bottom, the tertiary structure of msDNA. (**D**) Secondary structure of Ec86 msDNA based on the solved structure.

**Fig. S6.**
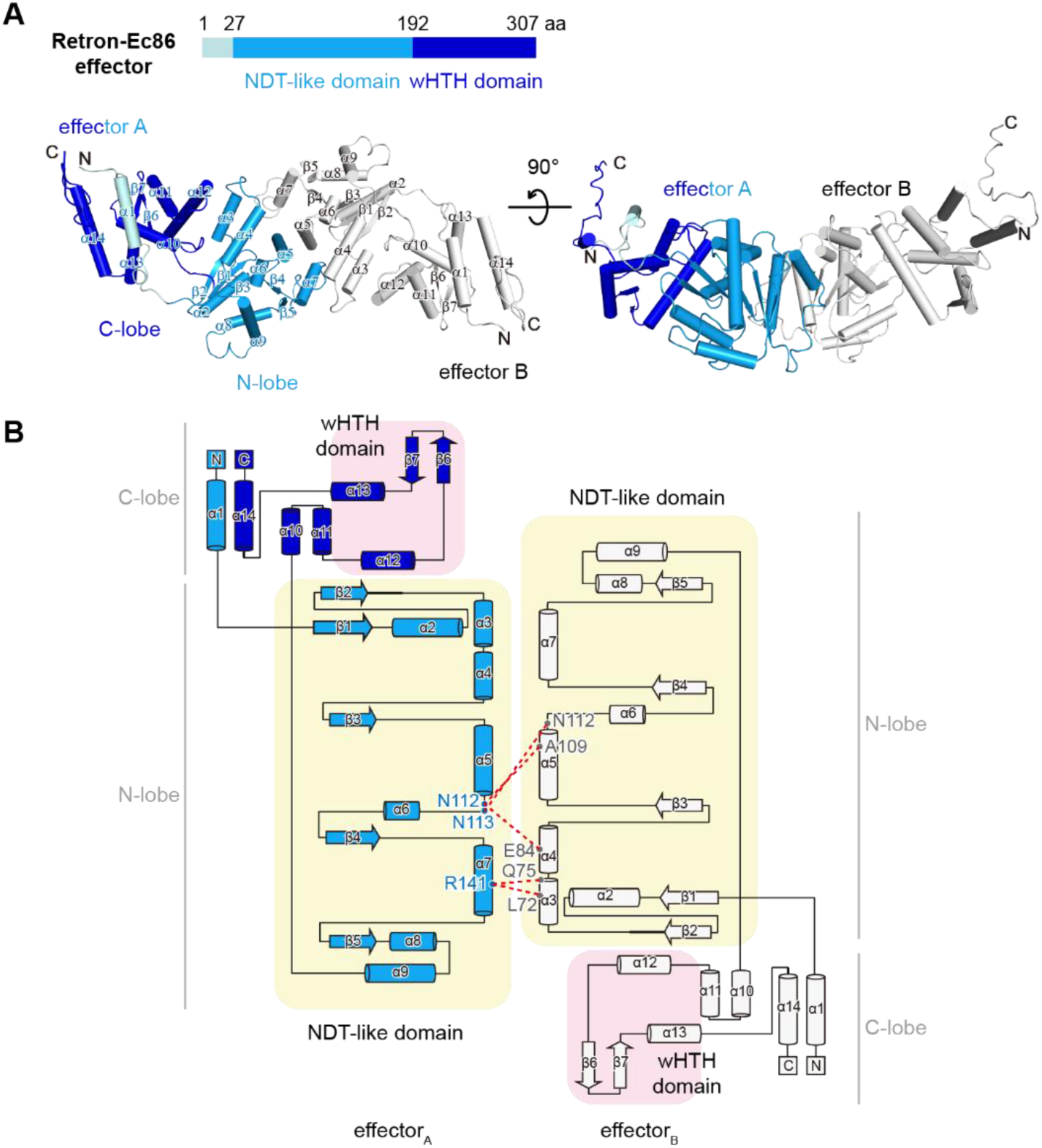
Ec86 effector dimer in a repeating unit of the Ec86-effector filament. (**A**) Schematic representation showing the domain architectures of the Ec86 effector and its atomic model in matching colours. α-helices and β-strands are labelled. N and C denote the N and C termini of the protein. α1 in effector_A_, palecyan; NDT domain in effector_A_, marine; winged HTH domain in effector_A_, blue; effector_B_, white. (**B**) Schematic diagram of Ec86 effector dimer. The NDT domain and wHTH domain of Ec86-effector are shaded in yellow and pink, respectively. The interacting residues in dimer interface are labeled and connected by red dotted lines.

**Fig. S7.**
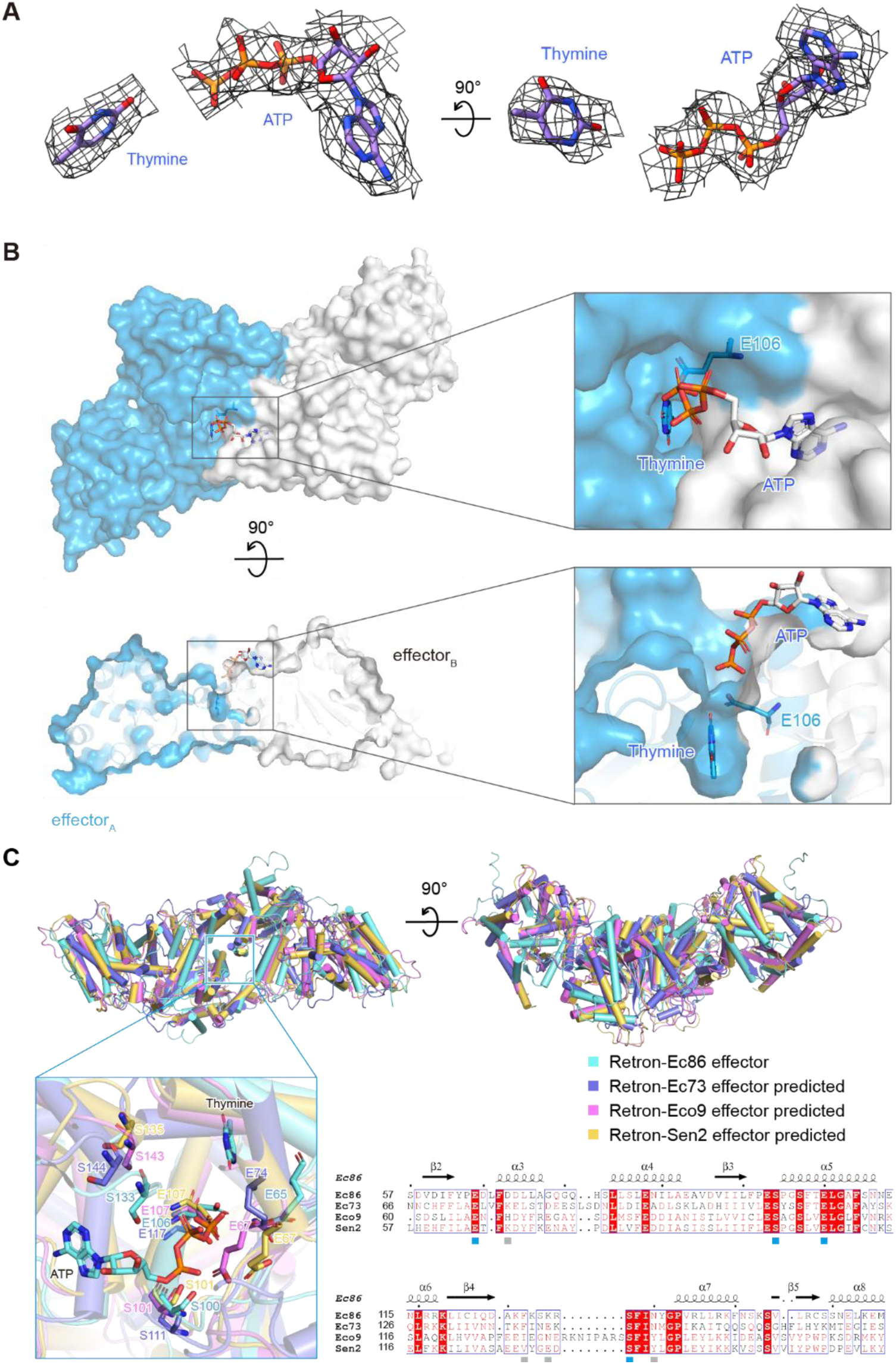
Structural analysis of the effector active-site_A_. (**A**) Cryo-EM density of the thymine and ATP. (**B**) Right, the ligand-binding pockets of Ec86-effector and in electrostatic surface potential. Left, the close-up view of active-siteA. The ligands and catalytic residue (E106) were shown as sticks and labeled. (**C**) Superposition of the NDT-like effectors from different retrons. The effectors of Ec73, Eco9 and Sen2 were predicted by alphafold2. Top panel, superpositions of Ec86-effector with Ec73-effector (RMSD=5.76), Eco9-effector (RMSD=6.03), and Sen2-effector (RMSD=6.47). Bottom panel, the close-up view of conserved residues interacting with ligands and the sequence alignment of corresponding region. Ec86 effector, cyan; Ec73 effctor, slate; Eco9 effctor, violet; Sen2 effctor, yelloworange.

**Fig. S8.**
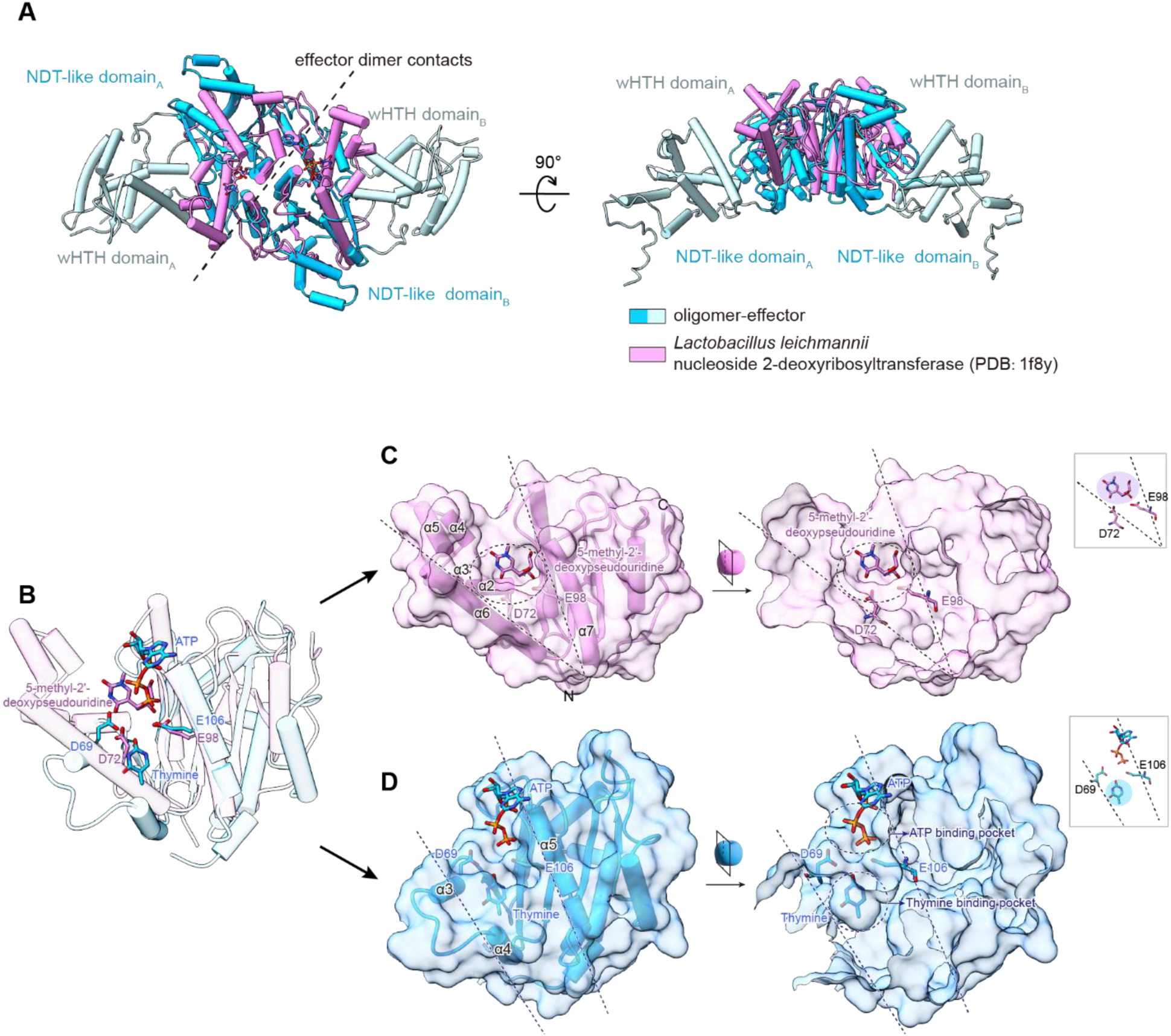
Comparisons of Ec86 effector with a classic NDT. (**A**) Superposition of the Ec86 effector and *Ll*NDT (pDB: 1f8y, RMSD of 4.99 over 128 residues). C-lobe in Ec86 effector, light blue; N-lobe in Ec86 effector, deep sky blue; *Ll*NDT, orchid. (**B**) Superposition of *Ll*NDT and the NDT-like domain of Ec86 effector. The ligands in active-site were shown as sticks and labeled. (**C**-**D**) surface representions of the ligand-binding pockets of Ec86 effector NDT-like domain (**C**) and *Ll*NDT (**D**).

**Fig. S9.**
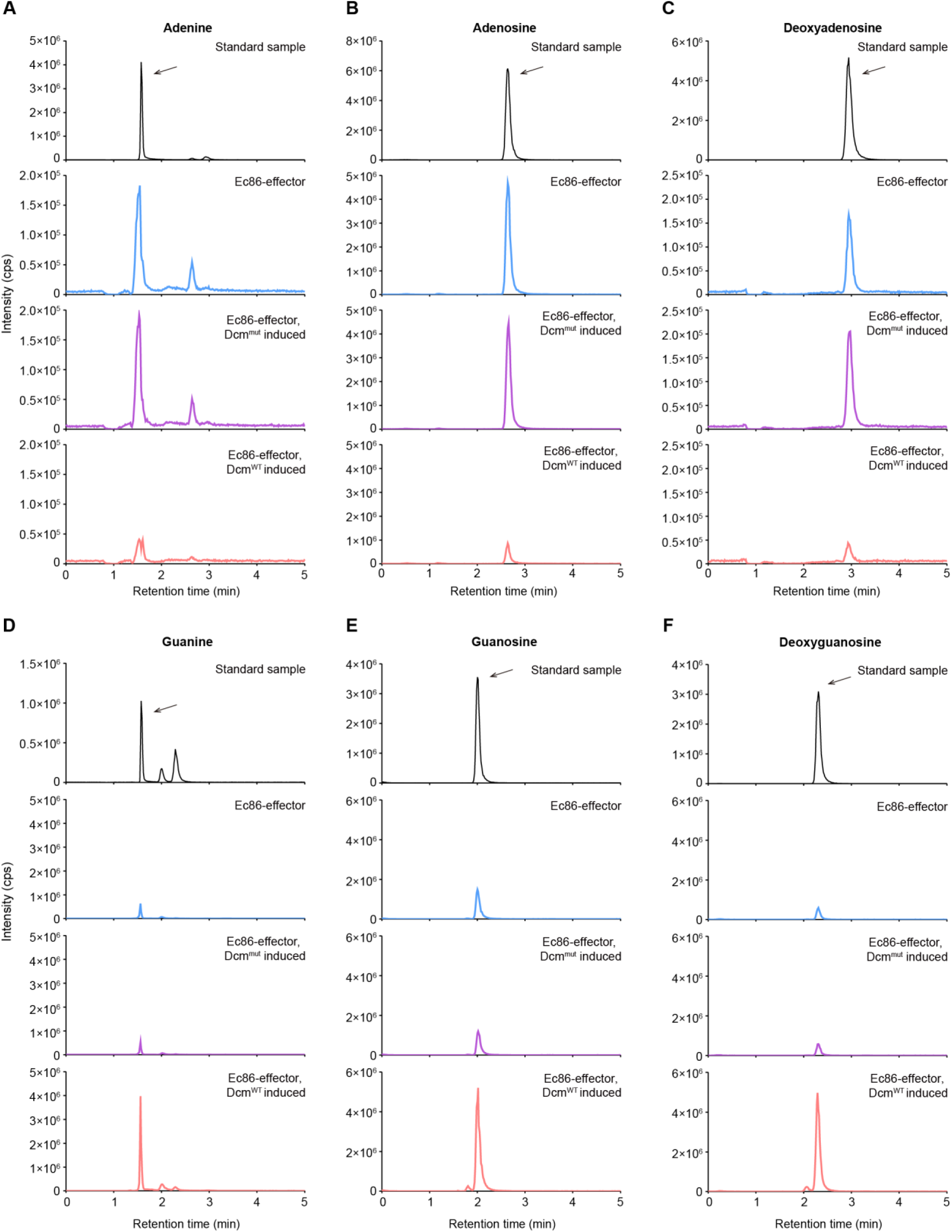
LC-MS analysis of the abundance of adenine and adenine derivatives (A-C) and guanine and guanine derivatives (D-F) in vivo. (**A-F**) Top panels, the adenine, adenosine, deoxyadenosine, guanine, guanosine and deoxyguanosine standards were detected using LC-MS. The in vivo abundance of these nucleosides and nucleoside derivatives in *E. coli* MG1655 cells expressing the retron Ec86 system from its native promoter after induction of Dcm expression, Dcm^mut^ (that contains C82A, E128A, K175A, and K176A mutations) expression, or a corresponding empty vector.

**Fig. S10.**
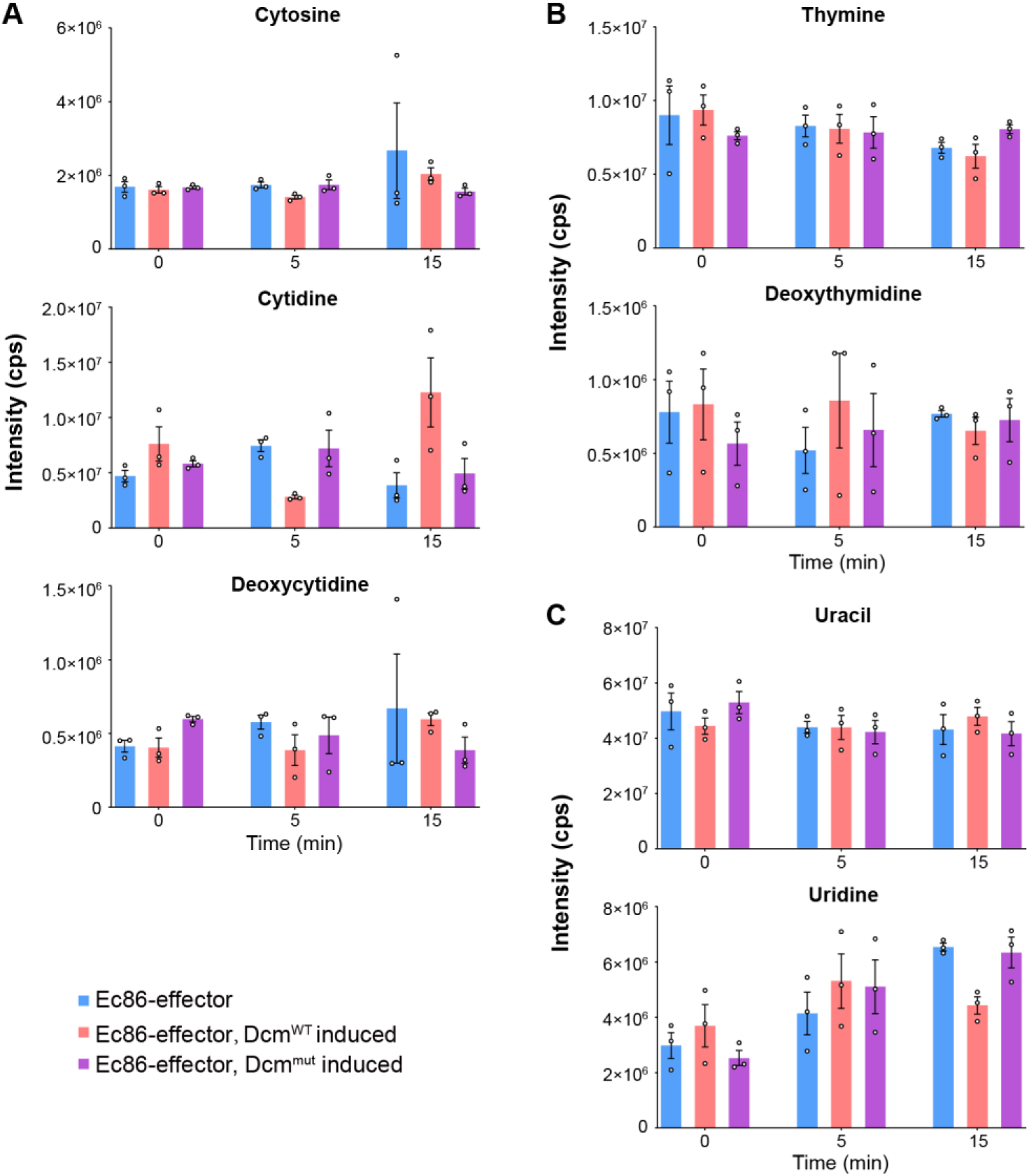
The activity of Ec86-effector defense system in vivo. (**A**-**C**) In vivo abundance of cytosine and cytosine derivatives (**A**), thymine and thymine derivative (**B**), and uracil and uracil derivative (**C**) in *E. coli* MG1655 cells expressing the retron Ec86 system from its native promoter after induction of Dcm expression, Dcm^mut^ (that contains C82A, E128A, K175A, and K176A mutations) expression, or a corresponding empty vector. Data represent the mean ± SEM of three biological replicate cultures.

**Table S1.**
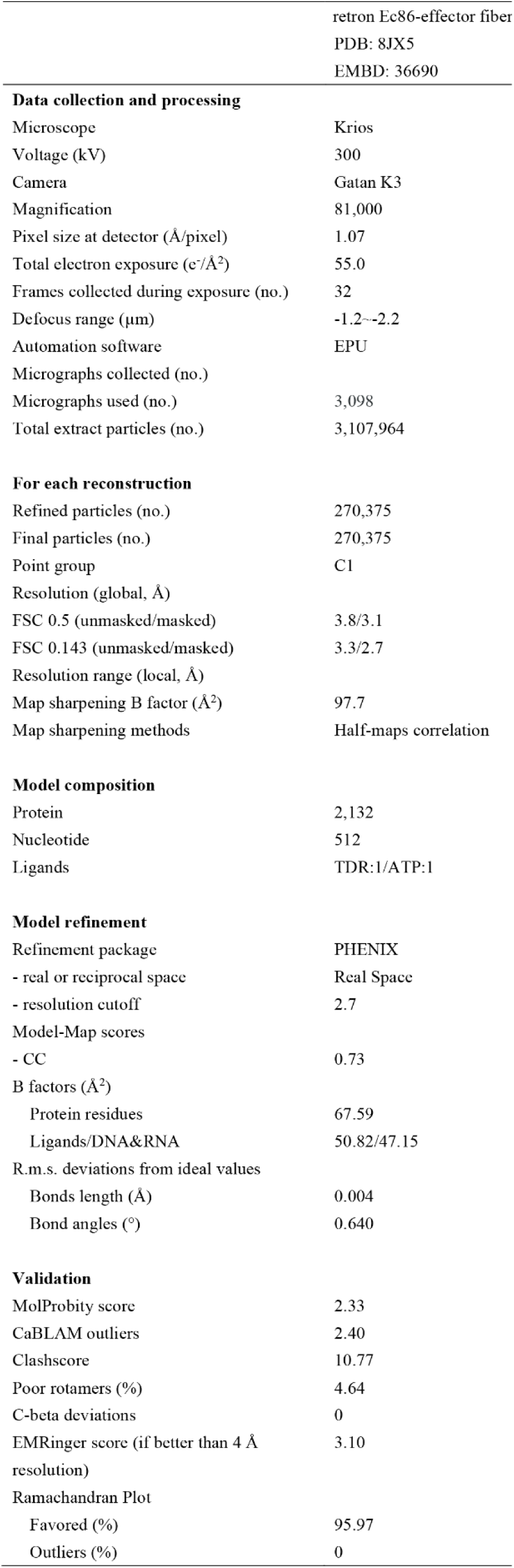
Cryo-EM data collection and refinement statistics.

|  |  |
| --- | --- |
|  | retron Ec86-effector fiber |
|  | PDB: 8JX5 |
|  | EMBD: 36690 |
| **Data collection and processing** |  |
| Microscope | Krios |
| Voltage (kV) | 300 |
| Camera | Gatan K3 |
| Magnification | 81,000 |
| Pixel size at detector (Å/pixel) | 1.07 |
| Total electron exposure (e<sup>-</sup>/Å<sup>2</sup>) | 55.0 |
| Frames collected during exposure (no.) | 32 |
| Defocus range (µm) | -1.2~-2.2 |
| Automation software | EPU |
| Micrographs collected (no.) |  |
| Micrographs used (no.) | 3,098 |
| Total extract particles (no.) | 3,107,964 |
| **For each reconstruction** |  |
| Refined particles (no.) | 270,375 |
| Final particles (no.) | 270,375 |
| Point group | C1 |
| Resolution (global, Å) |  |
| FSC 0.5 (unmasked/masked) | 3.8/3.1 |
| FSC 0.143 (unmasked/masked) | 3.3/2.7 |
| Resolution range (local, Å) |  |
| Map sharpening B factor (Å<sup>2</sup>) | 97.7 |
| Map sharpening methods | Half-maps correlation |
| **Model composition** |  |
| Protein | 2,132 |
| Nucleotide | 512 |
| Ligands | TDR:1/ATP:1 |
| **Model refinement** |  |
| Refinement package | PHENIX |
| - real or reciprocal space | Real Space |
| - resolution cutoff | 2.7 |
| Model-Map scores |  |
| - CC | 0.73 |
| B factors (Å<sup>2</sup>) |  |
| Protein residues | 67.59 |
| Ligands/DNA&RNA | 50.82/47.15 |
| R.m.s. deviations from ideal values |  |
| Bonds length (Å) | 0.004 |
| Bond angles (°) | 0.640 |
| **Validation** |  |
| MolProbity score | 2.33 |
| CaBLAM outliers | 2.40 |
| Clashscore | 10.77 |
| Poor rotamers (%) | 4.64 |
| C-beta deviations | 0 |
| EMRinger score (if better than 4 Å resolution) | 3.10 |
| Ramachandran Plot |  |
| Favored (%) | 95.97 |
| Outliers (%) | 0 |

